# Generalisable tissue-wide molecular reconstruction from histology

**DOI:** 10.64898/2026.06.09.731252

**Authors:** Andrew Zhang, Lijia Yu, Beilei Bian, Yue Cao, Shuchang Ye, Ethan Han, Harry Robertson, Yuhao Dong, Yixiao Mao, Boxiang Liu, Ellis Patrick, Jinman Kim, Jean YH Yang

## Abstract

Spatial transcriptomics technologies measure gene expression within intact tissues but remain difficult to scale across large tissue sections and patient cohorts. Consequently, many studies rely on tissue microarrays (TMAs) or sparse spatial profiling designs, where molecular measurements are available for only limited tissue regions and are often generated using heterogeneous gene panels.

Existing H&E to spatial gene expression prediction methods remain challenged by sparse molecular measurements, partially overlapping gene panels and tissue-wide reconstruction across heterogeneous spatial datasets. Here, we present GHIST+, a framework for tissue-wide reconstruction of single-cell molecular states from H&E histology. GHIST+ integrates cellular morphology, local tissue context and shared tissue representations to extend sparse molecular measurements into tissue-wide molecular maps across heterogeneous spatial datasets. Across multiple cancer types and GTEx breast tissues, GHIST+ reconstructs biologically meaningful tissue-wide molecular organisation from sparse TMA-derived measurements while preserving spatial tissue structure, cell-type organisation and age-associated tissue states across cancer and non-cancer settings. GHIST+ establishes a scalable framework for transforming sparse spatial profiling experiments into tissue-wide molecular maps, enabling cohort-scale molecular reconstruction from routine histology under heterogeneous spatial transcriptomic settings.

## Introduction

The advent of spatial omics technologies has transformed tissue biology by enabling molecular profiling within intact tissue architecture. However, spatial transcriptomic (ST) profiling remains expensive, whereas H&E histology is routinely acquired across both research and clinical settings. This difference has driven the development of deep learning approaches for H&E-to-SGE prediction that learn shared morphological-molecular representations from paired H&E images and spatial transcriptomic data to infer spatial gene expression (SGE). Early approaches were primarily developed for spot-based ST platforms^1^, enabling prediction of molecular states at multicellular spot resolution^1–4^. More recent studies have leveraged emerging single-cell and subcellular spatial technologies to enable single-cell-resolution reconstruction of gene expression and cellular states ^5–7^. Collectively, these approaches highlight the potential to extend molecular reconstruction across large histopathology repositories using routinely acquired H&E images. As spatial transcriptomics becomes increasingly adopted in clinical research, WSI profiling across large patient cohorts remains prohibitively expensive, motivating practical strategies for generating cohort-scale spatial molecular data.

Tissue microarrays (TMAs) provide a scalable and cost-effective strategy for high-throughput molecular profiling by enabling simultaneous profiling of representative tissue cores from multiple samples on a single slide. While this design substantially reduces experimental cost and improves cohort scalability, tissue cores in TMAs typically capture only a small fraction (sometimes <10%) of the original WSI, often missing important spatial information, including tumour microenvironment structure, spatial niches, and regional heterogeneity^8,9^. However, each tissue core in TMA has a corresponding WSI, creating an opportunity to computationally propagate molecular information from sparsely profiled tissue cores to the surrounding tissue landscape. This creates a new direction for H&E-to-SGE prediction, moving beyond molecular prediction from histology alone toward tissue-wide molecular reconstruction from sparse spatial measurement, a process we term “core-to-WSI”. The increasing availability of TMA-based and multi-cohort spatial datasets, often characterised by differing gene panels, tissue contexts, and experimental platforms, further highlights both the opportunity and the methodological challenges of tissue-wide molecular reconstruction from sparse spatial measurements.

These emerging data settings introduce several technical challenges for H&E-to-SGE modelling. In practice, models must generalise across cohorts, accommodate limited overlap between gene panels, recover unmeasured molecular features, and infer tissue-wide molecular states from sparsely profiled regions. Although recent advances in single-cell and subcellular spatial transcriptomics have accelerated the development of molecular reconstruction methods, many existing approaches were not designed for these settings. Existing approaches have largely addressed modelling improvements, such as feature extraction or spatial representation learning. For example, some approaches enhance histological feature extraction through pretrained pathology foundation models^6,7^, others use multimodal alignment between histology and gene-expression features^7^, while additional methods incorporate spatial context through graph-based modelling^6^ or introduce cell-type-aware supervision for biologically meaningful single-cell molecular reconstruction^5^. However, these advances leave behind the broader problem of tissue-wide reconstruction under sparse (e.g., with TMA tissue regions) and heterogeneous molecular measurements (e.g., different cancer) unresolved. These limitations constrain the ability of current methods to transform sparse spatial profiling experiments into tissue-wide molecular maps across heterogeneous cohorts.

Addressing this gap requires tissue-wide reconstruction architectures capable of propagating information across local cellular neighbourhoods and shared tissue states, while accommodating sparse and partially overlapping gene measurements across datasets.

To this end, we developed GHIST+, a framework for tissue-wide reconstruction of single-cell gene expression and cell identity from H&E histology under sparse spatial profiling, limited overlap between gene panels, and heterogeneous multi-task training settings. In TMA settings, GHIST+ uses sparse molecularly profiled tissue cores as *anchors* for reconstructing molecular landscapes across paired WSI images. We achieved this through three key innovations each targeting a distinct constraint of tissue-wide reconstruction from sparse spatial measurements. First, we introduce an Edge-Conditioned Residual Mixer (ECRM) to reconstruct cell-level molecular states beyond profiled cores by incorporating local cellular neighbourhood context. Second, we propose Vector Quantisation (VQ)-based tissue prototyping to address sparse spatial profiling by learning a compact set of recurring tissue states across the WSI. Third, we introduce a Mask-Aware Gene Conditioned Imputer (MAGCI) to address heterogeneous gene-panel coverage by learning from observed genes while imputing genes absent from a given panel. Across multiple cancers and tissue types, including non-cancerous tissues, GHIST+ enables accurate molecular reconstruction from sparse spatial measurements and supports biologically meaningful analysis across tissue contexts and biological states. The GHIST+ framework increases the utility of sparse spatial profiling experiments by enabling limited TMA-derived molecular measurements to be extended into tissue-wide molecular maps and integrated across heterogeneous spatial datasets, positioning GHIST+ as a generalisable framework for studying tissue organisation and dynamics.

## Results

### GHIST+ framework for cell-resolved molecular reconstruction across heterogeneous slides

GHIST+ is a generalisable framework for reconstructing cell-resolved molecular states from routine H&E histology across slides, gene panels and tissue types (**Fig. 1**). It is trained on paired H&E images and single-cell-resolution spatial data (**Fig. 1a**), and is designed for core-to-WSI reconstruction, where sparsely profiled tissue cores are used to infer cell-level molecular states across unprofiled WSI regions. This setting couples three challenges: cell states beyond profiled cores depend on local tissue context, TMA cores sample only sparse regions of the WSI, and gene panels differ across datasets. GHIST+ addresses these challenges through three linked modules. It first partitions paired images into fixed-size tiles and uses a pathology foundation encoder (UNI2h)^10^ to extract cell-level morphological features. ECRM then incorporates local cellular context by building an adaptive graph around each target cell, where edge features encode relative position, distance, morphology similarity and cell-type agreement determine which neighbouring cells refine the target representation. VQ maps continuous tile features to learned tissue prototypes, allowing recurring tissue states to connect profiled cores with unprofiled WSI regions and support representative region selection. MAGCI uses an observed-gene mask during panel completion, excluding unavailable genes from the training target while learning to recover genes absent from a given panel. Together, these modules reconstruct a cell-by-gene matrix under partially observed molecular measurements by combining morphology, local neighbourhood context, recurring tissue states and available gene measurements within a common output space (**Fig. 1b**).

**Figure 1.**
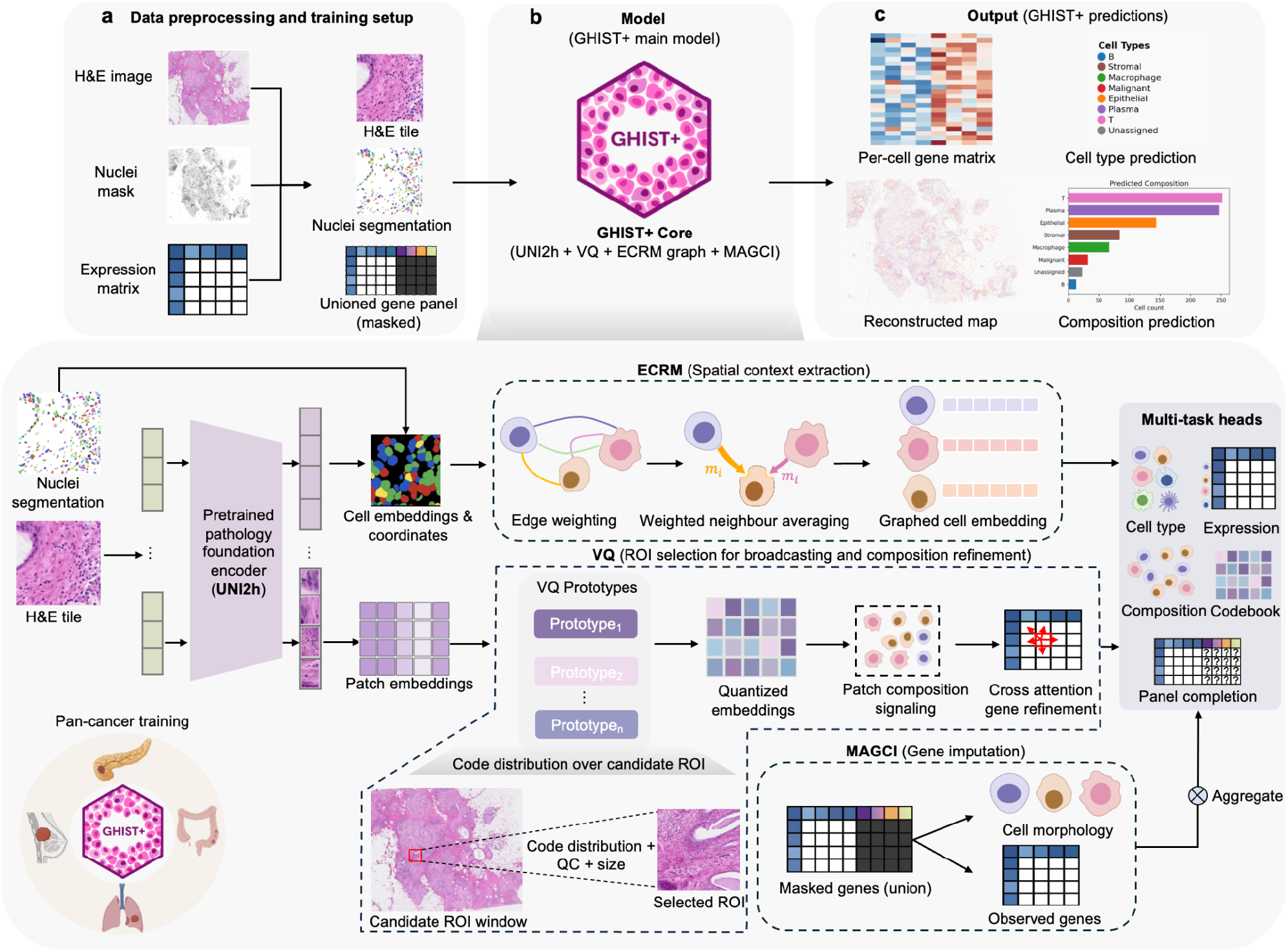
GHIST+ framework for molecular reconstruction from routine H&E histology. **a)Data preprocessing and training setup**. H&E images, nuclei segmentation masks and gene expression matrix are tiled and aligned under a shared gene panel supporting multi panel training. **b)Model**. GHIST+ maps H&E images to spatially resolved single-cell gene expressions through three key innovations: **Edge-conditioned Residual Mixer (ECRM)** that propagates cell information over a neighbourhood graph, **Vector Quantisation prototyping (VQ)** for patch composition signalling and TMA region selection and a **Mask-Aware Gene Conditioned (MAGCI)** for unified gene panel completion. c) **Outputs**. GHIST+ primarily outputs per-cell gene expression whereas cell types, segmentation and patch composition are used as auxiliary supervision in the multi task framework.

**Fig. 1c** shows that GHIST+ reconstructs cell-resolved molecular maps from H&E histology across both measured and unmeasured tissue regions. Ablation experiments showed that all three modules and the foundation encoder contribute to reconstruction performance (**Supplementary Fig. S1**).

Based on correlation on the top 20 spatially variable genes (SVGs), removing VQ, ECRM and the foundational pathology encoder reduced performance by 5.5%, 11.5% and 16.9%, respectively.

### GHIST+ improves tissue-wide molecular reconstruction using sparse TMA-derived molecular anchors

We first investigate whether sparse molecular measurements from TMA cores could serve as **molecular** anchors for core-to-WSI reconstruction. Within the BreastCancer2 dataset, visually similar histological regions exhibited distinct molecular programmes, including differences in immune, apoptosis and hormone-response signalling despite comparable desmoplastic morphology (**Fig. 2a**). For example, *IL2-STAT5* signalling showed stronger enrichment in the immune-infiltrated region B and desmoplastic region C relative to region A (**Fig. 2b**). Top 10 pathway rankings further demonstrated distinct regional transcriptional programmes, with regions B and C showing stronger enrichment of immune- and apoptosis-associated pathways, whereas region A showed relatively higher enrichment of epithelial and hormone-response-related programmes (**Fig. 2c**). These findings indicate that morphology alone may not sufficiently resolve tissue molecular states and motivate the use of molecular anchors to guide **core-to-WSI reconstruction**, which is also known as “broadcasting”^11^.

**Figure 2.**
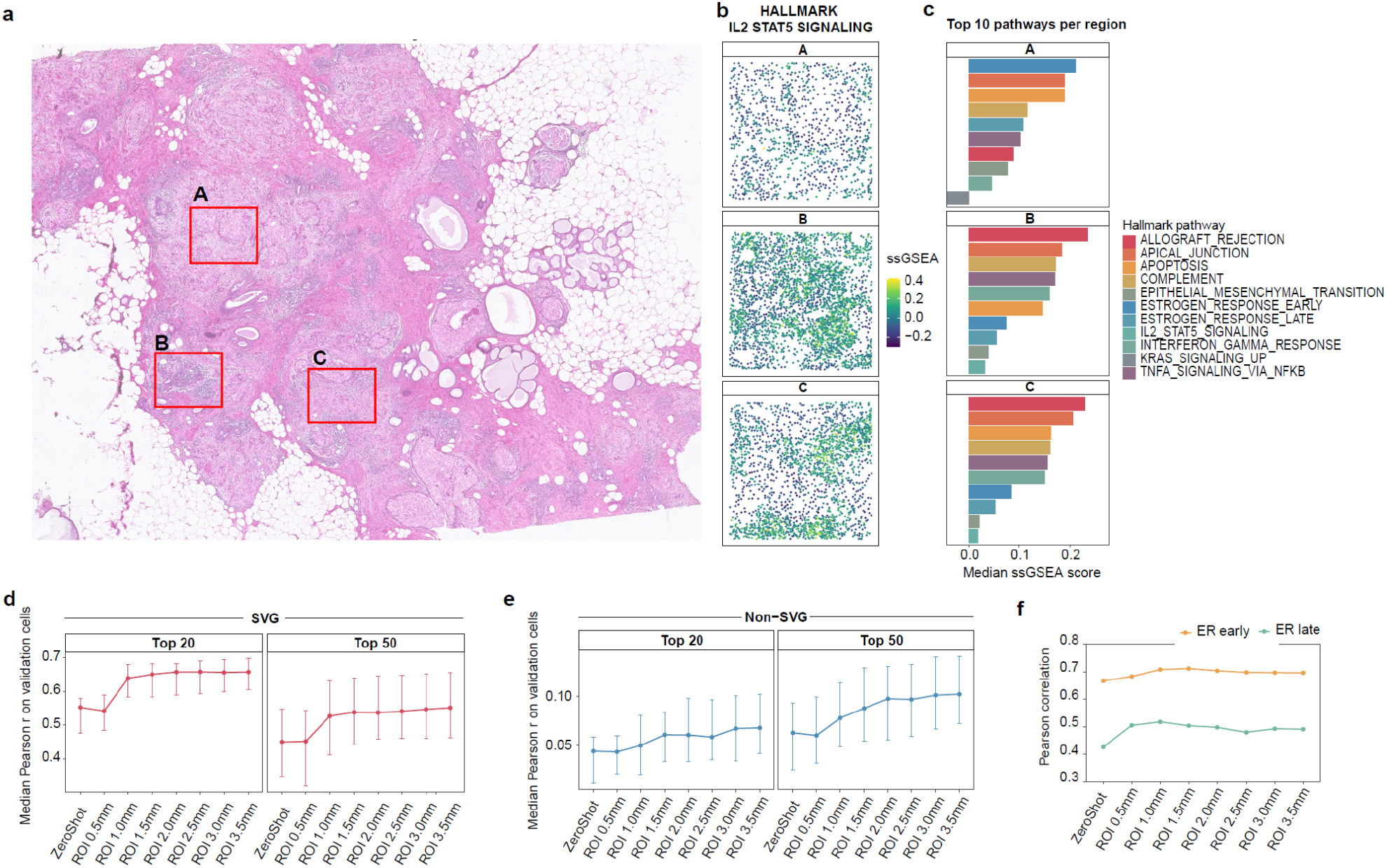
GHIST+ supports pathway-aware broadcasting from limited TMA-like calibrated regions. (a) Three selected regions in the BreastCancer2 slide, region A and C are desmoplastic tumour-associated stroma with embedded epithelial tumour nests, and region B is a densely immune-infiltrated microenvironment with lymphoid-like morphology. (b) Spatial distribution of ssGSEA scores for the *IL2-STAT5* signalling pathway across three selected regions. c) Top 10 enriched Hallmark pathways ranked by median ssGSEA score in each selected region. (d, e) Median Pearson correlation on validation cells in spatially held out regions for top 20 and top 50 SVGs (d) and non-SVGs (e) under different ROI sampling sizes (0.5-3.5mm^2^) and zero-shot inference settings. Error bars indicate variability across repeated evaluations. (f) Pearson correlation between ssGSEA scores derived from predicted and experimentally measured Xenium data for early estrogen response and late estrogen response signatures across ROI sampling sizes in TMA-anchor and zero-shot models.

To test whether molecular anchors improve core-to-WSI reconstruction, we compared GHIST+ predictions with and without profiled TMA cores. In the anchored setting, molecular measurements from selected TMA cores were provided to the model, and performance was evaluated on withheld regions from the same WSI. This design tests whether sparse local molecular measurements improve reconstruction beyond H&E-only inference. We selected TMA cores from VQ-defined tissue areas and progressively increased the profiled core size from 0.5 mm^2^ to 3.5 mm^2^ to assess how reconstruction performance changes with increasing tissue coverage (**Supplementary Fig. S2**). Core measurements were then provided to a pretrained GHIST+ model to reconstruct gene expression in withheld tissue regions. Increasing anchors’ size consistently improved prediction accuracy for spatially variable genes, with Pearson correlation increasing by 14.5% and 10% for the top 20 and 50 SVGs, respectively (**Fig. 2d, e**). Larger TMA core anchors generally yielded improved reconstruction performance, representing a trade-off between molecular profiling area and predictive accuracy. When TMA core anchors were provided, GHIST+ showed improved spatial concordance with ground-truth pathway maps compared with H&E-only reconstruction.

Pearson correlation for the early estrogen-response signature increased from 0.644 to 0.691 using a 1.5 mm^2^ core anchor, whereas the late estrogen-response signature improved from 0.435 to 0.522 using a 1 mm^2^ core anchor (**Fig. 2f; Supplementary Fig. S3**).

### GHIST+ preserves molecular and spatial tissue structure across-slides

We compared GHIST+ against GHIST^5^ and SpatialEx^6^ using the Xenium BreastCancer2 dataset under both within-slide and independent-slide evaluation settings. GHIST+ achieved the strongest performance across spatially variable and highly variable genes, with improved preservation of molecular and spatial tissue structure relative to existing methods (**Supplementary Fig S4**). In within-slide evaluation, GHIST+ achieved the highest median PCC across the top 20 SVGs (0.744 versus 0.714 and 0.521 for GHIST and SpatialEx, respectively; **Fig. 3a)**. These improvements remained consistent in independent-slide evaluation, where GHIST+ retained substantially stronger predictive performance across unseen tissue sections (median PCC = 0.653; **Fig. 3c, Supplementary Fig S5**), while also preserving higher-order gene co-expression structure as measured by Correlation Matrix Distance (CMD)(**Fig. 3d)**.

**Figure 3.**
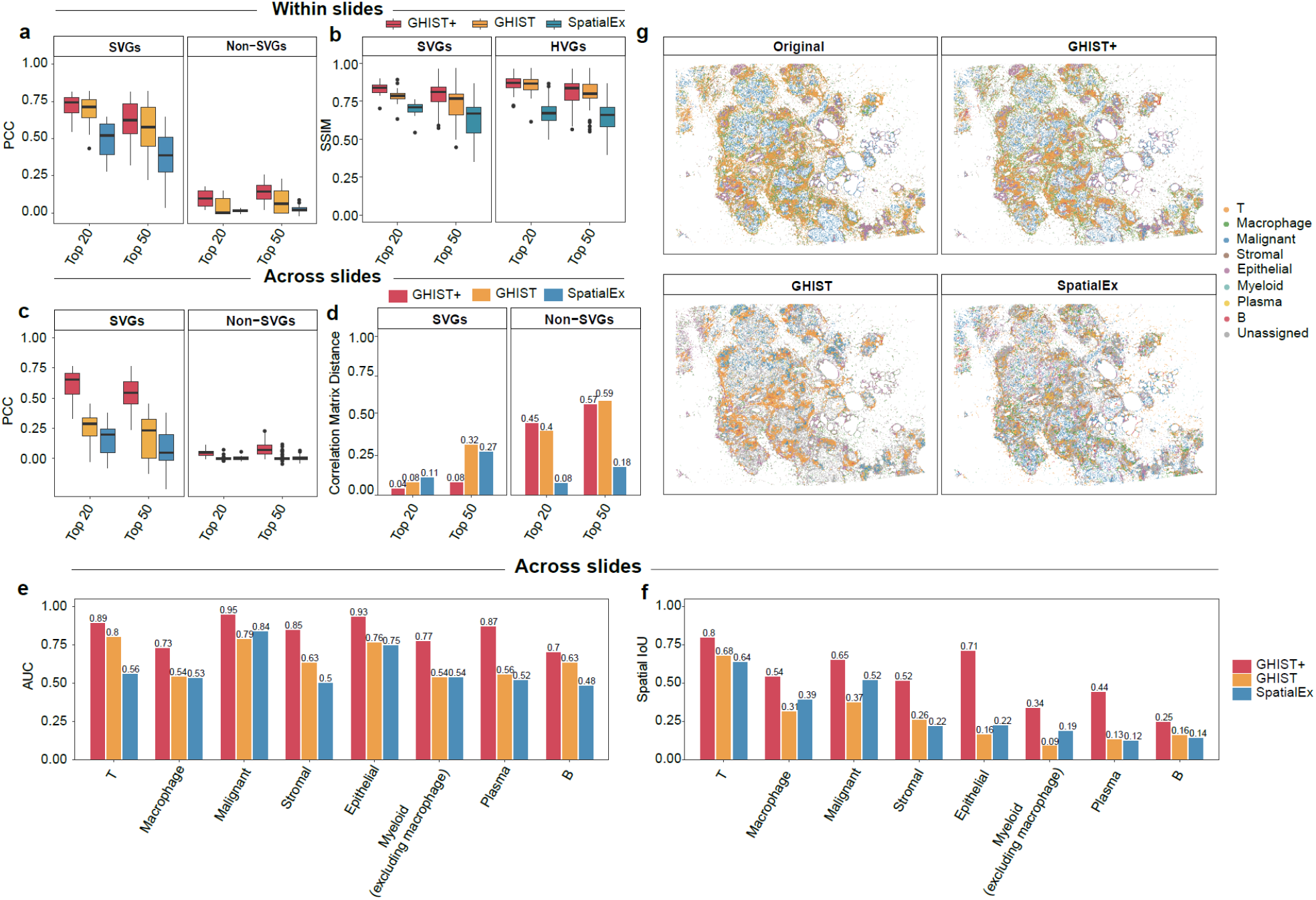
GHIST+ improves within-slide and across-slide cell-level expression prediction compared to state-of-the-art. (a,b) Boxplot comparison between GHIST+ with state-of-the-art subcellular gene expression prediction methods within-slides measured using per gene PCC and SSIM on the top 20 and top 50 SVGs and non-SVGs. (c) Boxplot comparison of per-gene PCC across slides for GHIST+ and existing subcellular gene expression prediction methods using top 20 and top 50 SVGs and non-SVGs. (d) Barplot comparing Correlation Matrix Distance between predicted and ground truth gene-gene correlation structure across all genes and top 20 and 50 SVGs. (e) Barplot measuring across-slide recovery of cell-types inferred from predicted gene expression, quantified by AUC on eight cell types (T, Malignant, Epithelial, Myeloid, Plasma, Macrophage, Stromal, B). (f) Barplot comparing per class spatial intersection of individual cell types with IoU on the same eight cell types. g) Cell type annotation maps reconstructed from gene markers and compared with ground truth.

GHIST+ also more accurately reconstructed cell-type organisation and tissue architecture. Predicted cell-type distributions more closely matched observed spatial organisation, with improved recovery of malignant and immune cell regions relative to existing methods (Supplementary **Fig. S4d, e**).

Across both within-slide and independent-slide evaluations, GHIST+ achieved stronger preservation of spatial structure and tissue compartmentalisation, including higher SSIM, IoU and AUROC scores (**Fig. 3b, e; Supplementary Fig. S6**). Together, these findings indicate that GHIST+ improves reconstruction accuracy and preserves biologically meaningful molecular and spatial patterns across tissue sections.

### GHIST+ reconstructs unmeasured genes across heterogeneous spatial transcriptomic panels

Since gene panel overlap across single-cell resolution ST datasets is limited, particularly across cancer types, models must impute unmeasured genes during inference on unseen samples. We therefore evaluated GHIST+ against expression only imputation methods PASTA^12^, SpatialEx, SpaGE^13^, SpaIM^14^, stImpute^15^ and Tangram^16^ across five Xenium breast slides using synthetically masked genes. GHIST+ achieved the strongest overall imputation performance, including the highest median PCC (0.743, 0.747) and lowest median RMSE (2.904, 3.010) across held-out genes (**Fig. 4a; Supplementary Fig. S7**), while also preserving higher-order gene co-expression structure with consistently low CMD across-slides (CMD < 0.15; **Fig. 4b**). GHIST+ achieved the lowest predictive error, as measured by a TISSUE calibration score of 0.343^17^, compared to competing methods (**Fig. 4a**).

**Figure 4.**
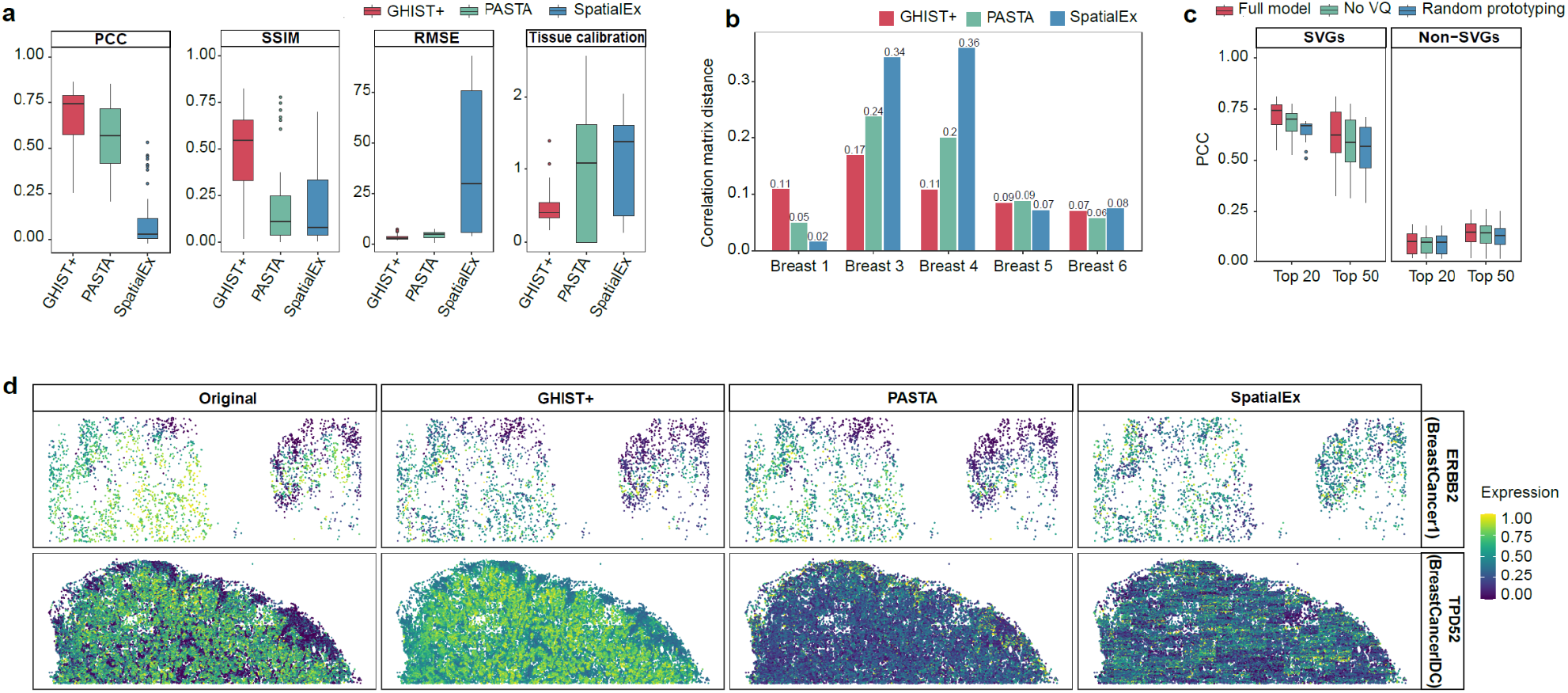
GHIST+ accurately imputes held-out genes and benefits from learned prototyping. (a) Boxplot comparison of imputation performance between state-of-the-art cell-level imputation methods measured using per gene PCC, SSIM, RMSE and TISSUE calibration score. (b) Barplot comparing GHIST+ imputation performance per slide on holdout genes against state-of-the-art cell-level imputation methods measured using Correlation Matrix Distance. (c) Boxplot showing VQ ablation study comparing full model, no VQ variant and random prototyping evaluated with per gene PCC on the top 20 and top 50 SVGs and HVGs. (d) Representative spatial expression maps for selected holdout genes (ERBB2 and TPD52).

Beyond expression recovery, GHIST+ more faithfully preserved spatial gene organisation. Across held-out genes, GHIST+ achieved the highest SSIM scores (**Fig. 4a**), with predicted spatial patterns for representative genes such as *ERBB2* and *TPD52* closely matching ground-truth tissue structure (**Fig. 4d**). Ablation analyses showed that removing learned VQ prototypes reduced reconstruction performance by ∼5-6%, while replacing them with random prototypes resulted in performance drops of up to ∼10% for top SVGs (**Fig. 4c**), demonstrating that tissue-level prototypical context contributes substantially to unmeasured gene recovery beyond local cell morphology alone.

### Pan-cancer training improves generalisability across tissue contexts

Spatial transcriptomics remains costly, resulting in limited sample sizes for many different cancers, particularly for rare or heterogeneous ones. To address this, GHIST+ adopts a pan-cancer training strategy for domain generalisation (**Supplementary Fig. S8**). By learning a shared H&E -to-gene latent space across tumour types, the model can use information from other cancers to improve prediction when only limited training data are available. We compared two scenarios for prediction on PancreasADC3: (i) single-slide models trained on PancreasADC1 or PancreasADC2, and (ii) pan-cancer-assisted zero-shot inference. Single slide models transferred poorly. Pan-cancer assisted inference improved this transfer for both scenarios. Relative to the PancreasADC2 trained model, median PCC increased from 0.003 to 0.102 for the top 20 SVGs and from 0.003 to 0.172 for the top 50 SVGs. Relative to the PancreasADC1 trained model, median PCC increased from -0.001 to 0.236 for the top 20 SVGs (**Fig. 5a**). These results indicate that training across multiple cancer slides improves cross-slide gene expression prediction compared with models trained on a single slide.

**Figure 5.**
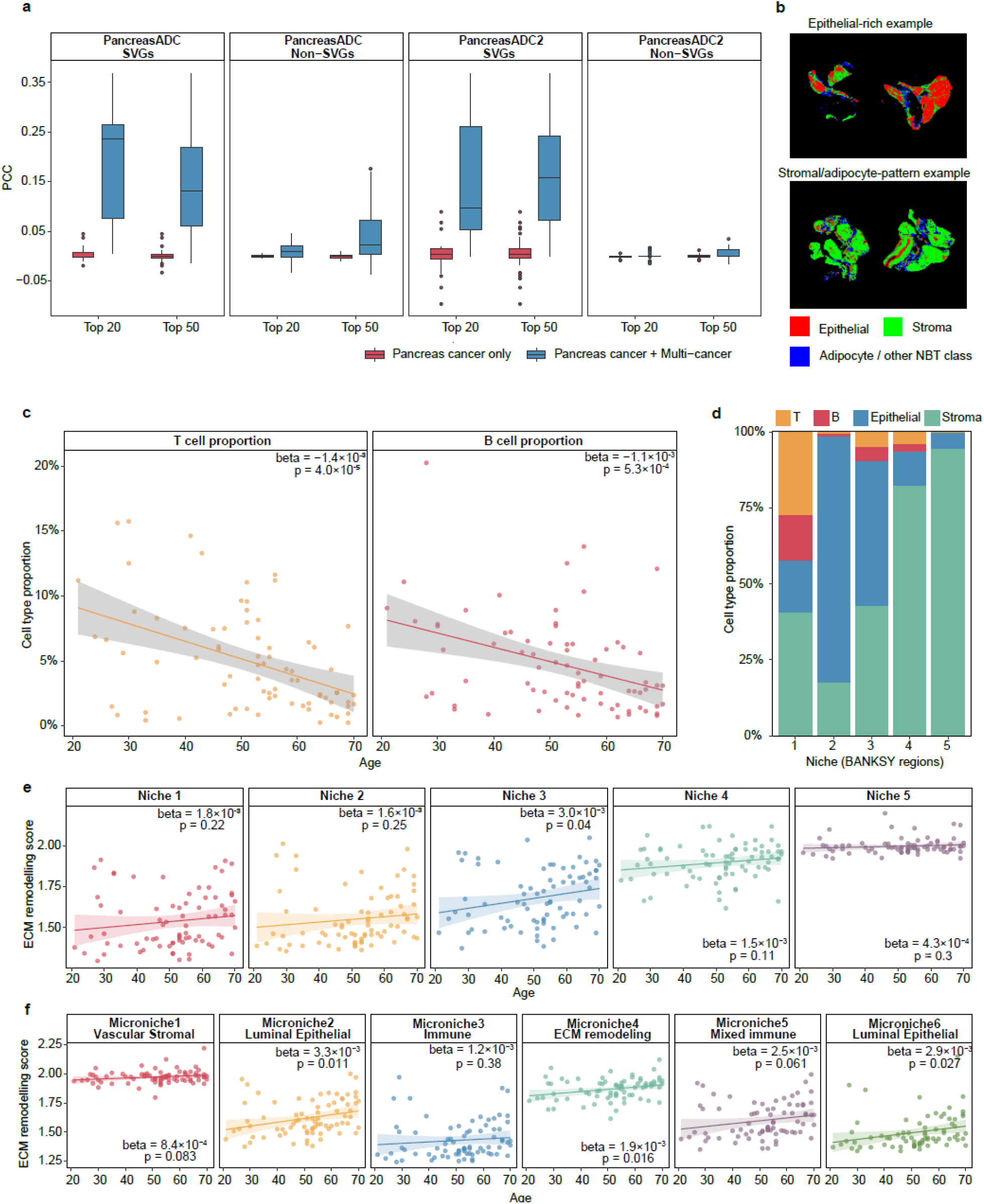
GHIST+ pan-cancer modelling improves single slide performance and generalizes from tumour slides to normal tissues. (a) Comparison of cross-slide generalisation measured by PCC for pancreas single-slide training and pan-cancer-assisted inference across top SVGs and non-SVGs. (b) Representative histology-guided tissue compartment maps from young (age 28; GTEX-15ER7-1626) and old (age 66; GTEX-1F88E-0326) GTEx normal breast samples. (c) Scatter plot of T and B cell type proportion within (y-axis) epithelial-immune niche (niche 3) against age (x-axis). Each point represents a sample. (d) Stacked bar plots show the cellular composition of major cell types within each spatial niche identified by BANKSY across all samples. (e) Scatter plot of ECM remodelling scores (y-axis) within each BANKSY niche against age (x-axis) in GTEx normal breast tissue. (f) Scatter plot of ECM remodelling scores (y-axis) against age for each of the microniches from the histology-defined epithelial compartments.

### GHIST+ generalises from tumour training to normal breast tissues and recovers age-associated spatial programmes

We assessed whether GHIST+ generalises beyond tumour settings using 77 normal GTEx female breast tissues (**Supplementary Fig. S9**). To characterise tissue organisation, we applied a supervised histology-guided classifier^18^ that identified epithelial, stromal, and adipose compartments (**Fig. 5b**). Marker analysis identified expected markers including KRT7 and TACSTD2 (**Supplementary Table 1**), supporting the ability of GHIST+ to reconstruct biologically meaningful spatial tissue architecture beyond tumour settings.

**Table 1.** Complete list of datasets used in the training of GHIST+. Tissues include Pancreas, Lung, Breast, Colorectal. Overall, there are 2,785,236 detected cells and 1,275 unique genes detected after preprocessing.

| Dataset name | Tissue | Detected Cells | Number of genes | ID in GitHub code | Description |
| --- | --- | --- | --- | --- | --- |
| BreastCancer1 | Breast | 94,000 | 313 | Breast 1 | High resolution mapping of the breast cancer tumor microenvironment using integrated single cell, spatial and in situ analysis of FFPE tissue (Sample 1) |
| BreastCancer2 | Breast | 86,344 | 280 | Breast 2 | High resolution mapping of the breast cancer tumor microenvironment using integrated single cell, spatial and in situ analysis of FFPE tissue (Sample 2) |
| BreastCancerIDC | Breast | 487,041 | 380 | Breast 3 | Xenium FFPE Human Breast with Custom Add-on Panel |
| BreastCancerIDC2 | Breast | 453,436 | 280 | Breast 4 | FFPE Human Breast with Pre-designed Panel (Sample 1) |
| BreastCancerILC | Breast | 311,000 | 280 | Breast 5 | FFPE Human Breast with Pre-designed Panel (Sample 2) |
| BreastCancerIDC3 | Breast | 295,909 | 380 | Breast 6 | FFPE Human Breast with Custom Add-on Panel |
| ColorectalNormal | Colorectal | 83,422 | 480 | Colorectal 1 | Human Colon Preview Data (Xenium Human Colon Gene Expression Panel) (Non-diseased, pre-designed panel) |
| ColorectalADC | Colorectal | 103,228 | 425 | Colorectal 2 | Human Colon Preview Data (Xenium Human Colon Gene Expression Panel) (Cancer, pre-designed panel) |
| ColorectalADC2 | Colorectal | 145,103 | 325 | Colorectal 3 | FFPE Human Colorectal Cancer Data with Human Immuno-Oncology Profiling Panel and Custom Add-on |
| LungADC | Lung | 118,476 | 480 | Lung 1 | Preview Data: FFPE Human Lung Cancer with Xenium Multimodal Cell Segmentation |
| LungNSCLC | Lung | 105,016 | 377 | Lung 2 | FFPE Human Lung Cancer Data with Human Immuno-Oncology Profiling Panel and Custom Add-on |
| LungADC2 | Lung | 218,642 | 289 | Lung 3 | Xenium Human Lung Preview Data |
| PancreasADC | Pancreas | 24,117 | 380 | Pancreas 1 | FFPE Human Pancreas with Xenium Multimodal Cell Segmentation |
| PancreasADC2 | Pancreas | 149,731 | 474 | Pancreas 2 | FFPE Human Pancreatic Ductal Adenocarcinoma Data with Human Immuno-Oncology Profiling Panel |
| PancreasADC3 | Pancreas | 109,771 | 377 | Pancreas 3 | Pancreatic Cancer with Xenium Human Multi-Tissue and Cancer Panel |

Next, niche analysis with BANKSY^19^ identified five spatial niches (**Fig. 5d**) and given the established role of immune decline and extracellular matrix (ECM) remodelling during breast ageing and epithelial involution^20–22^, we next examined age-associated cellular and molecular changes across these niches. A mixed epithelial-immune niche (Niche 3) showed significant age-associated declines in both B-cell and T-cell abundance (both P < 0.001; **Fig. 5c**; **Supplementary Fig. S10**), suggesting disruption of epithelial-associated immune microenvironments. Significant increases in ECM-remodelling scores with age were also observed within Niche 3 (P = 0.04; **Fig. 5e**), consistent with previous reports of local stromal remodelling during breast ageing^23^. In contrast, stromal-dominant niches (Niches 4 and 5; **Fig. 5e**) showed no significant age-associated changes. Notably, although Niche 3 contained mixed populations, it arose predominantly from histology-defined epithelial compartment, suggesting unresolved multicellular heterogeneity. We therefore performed secondary niche analysis within the epithelial compartment.

Here, we delineated distinct luminal epithelial, immune-associated, vascular stromal, mixed immune, and ECM-remodelling programmes within histology-defined epithelial compartment (**Supplementary Table 2**). **Fig. 5f** shows age-associated ECM-remodelling signals were enriched specifically within luminal epithelial-associated (Microniche 2 and 6; P = 0.0114 and 0.0266) and ECM-remodelling programmes (Microniche 4; P = 0.0156). These findings suggest that breast ageing-associated remodelling is concentrated within luminal epithelial-associated microenvironments rather than representing a global tissue-wide process. These results demonstrate that GHIST+ can reconstruct spatially resolved molecular programmes from H&E histology and uncover previously unresolved multicellular heterogeneity.

## Discussion

Here, we present GHIST+, a generalisable framework for reconstructing tissue-wide molecular landscapes at single-cell resolution from H&E histology across gene panels, slides and tissue types. By integrating cellular morphology with neighbouring-cell context, GHIST+ enables sparse molecular measurements to be extended into biologically meaningful whole-slide molecular maps while preserving spatial tissue organisation and cell-type structure. In practical settings, including TMA-based studies, GHIST+ reconstructs molecular information across whole tissue sections using only limited spatial measurements. Across heterogeneous cohorts, GHIST+ supports transfer beyond tumour-specific settings, accurately recovering normal tissue states and preserving interpretable spatial niches and age-associated tissue structure in GTEx breast tissues. Together, these findings position GHIST+ as a generalisable framework for studying tissue organisation across biological states from sparse spatial measurements.

GHIST+ is distinctive to other models because it reframes expression prediction from a regression task^24^ to a reconstruction task. Prior work suggests that morphology alone is insufficient to determine molecular states since visually similar cells can contain varying expressions^25^. In a tissue, a cell’s molecular state is shaped by not only its morphology, but also its surrounding tissue microenvironment^26^. Compared with hypergraph-based spatial models that aggregate broader neighbourhood structure^6^, our ECRM selectively uses neighbouring information according to spatial relationships, allowing each cell to be interpreted together with its local neighbourhood while still preserving single-cell resolution. This design allows GHIST+ to disambiguate cells with similar morphology and reconstruct molecular patterns that isolated H&E features are unlikely to capture.

Additionally, GHIST+ distinguishes itself by treating unmeasured genes as a component of tissue-wide molecular reconstruction. Current single-cell spatial transcriptomic technologies typically profile targeted gene panels with sparse and partially observed molecular measurements with substantial technical noise^27–29^. This is particularly relevant for technologies such as Xenium, where targeted molecular profiling enables single-cell resolution but does not provide complete transcriptome coverage. Through MAGCI, GHIST+ enables heterogeneous gene panels to contribute within a shared reconstruction framework, increasing the usable molecular information obtained from each slide. This process is further supported by VQ prototyping, which represents tissue regions through recurring tissue states so that recovery of unmeasured genes is guided not only by observed expression, but also by shared tissue context. Consequently, each profiled region contributes both molecular and tissue-state information, supporting more robust cross-slide and pan-cancer molecular reconstruction from sparse spatial measurements.

The ability of GHIST+ to infer virtual transcriptomic landscapes from routine H&E slides has huge downstream impacts for precision medicine^30,31^. Recent studies such as S2Omics^11,32^ have explored optimisation of ROI selection for spatial omics experiments and TMA directly from H&E. However, as demonstrated by our core-to-WSI analyses, visually similar H&E regions can exhibit distinct molecular programmes, indicating that morphology alone may not provide sufficient information to guide TMA-core selection. In this context, GHIST+ provides a complementary capability by integrating transcriptomic inference into both TMA design and tissue-wide molecular reconstruction. GHIST+ predictions can guide ROI or TMA-core selection by identifying tissue regions enriched for clinically relevant cell states, immune niches or pathway activities that are not evident from morphology alone. Once sparse TMA cores are profiled, these regions can subsequently be used as anchors to propagate molecular information across the whole slide. By complementing experimental spatial omics assays, GHIST+ increases the value of limited molecular measurements and supports scalable tissue-wide molecular characterisation from routine histology slides for biomarker discovery and precision-medicine applications in large clinical cohorts.

A notable finding is that GHIST+ is not limited to pan-cancer modelling, but instead learns morphology-molecular relationships that generalise beyond disease-specific contexts. Pan-cancer training plays a critical role in enabling this, as it reduces dependence on slide-specific tumour programmes and encourages the model to capture patterns that are stable across heterogeneous tissues and partially overlapping gene panels. As such, this enables meaningful transfer to normal tissue, as demonstrated using data from the GTEx Consortium^33,34^, where predicted expression preserves interpretable spatial niches and age-associated structure^35^. More broadly, this adaptive training strategy may be particularly advantageous in data-limited settings, where transferable morphology-molecular representations can support robust molecular reconstruction across related tissue contexts. Given that comprehensive spatial transcriptomic profiling of normal tissues remains substantially less common than disease-focused cohorts, in silico reconstruction of normal tissue ecosystems may provide a valuable analytical reference for understanding disease-associated tissue remodelling and microenvironmental change. This shift from robustness across cancers to transferability across biological states positions GHIST+ as a generalisable framework for studying tissue organisation and dynamics, particularly in settings such as ageing where spatial omics data remain limited.

This transferability provides the foundation for advancing GHIST+ towards a ‘virtual cell’ framework, where molecular information at the level of individual cells can be reconstructed across WSI from routine H&E histology. By leveraging heterogeneous datasets, the GHIST+ pan-cancer training strategy supports modelling in data-limited settings and enables generalisation beyond tumour-specific contexts, including reconstruction of normal tissue states from tumour-trained models. GHIST+ is not intended to replace experimental measurements or perturbation studies; rather, it provides a computational approach that allows spatially resolved, tissue-wide predictions. In doing so, it supports hypothesis generation and enables more informed experimental design. In combination with emerging perturbation frameworks such as CPA^36^, GEARS^37^ and Celcomen^38^, GHIST+ may further support future modelling of spatially organised tissue ecosystems.

In summary, GHIST+ introduces a framework for tissue-wide molecular reconstruction from sparse and partially observed spatial transcriptomic measurements. By learning transferable histology-molecular representations across slides, gene panels and tissue types, GHIST+ extends sparse molecular profiling into biologically meaningful tissue-wide molecular maps across heterogeneous cohorts and biological contexts

## Supporting information

All supplementary figures

## Acknowledgements

The authors thank all their colleagues, particularly at The University of Sydney, Sydney Precision Data Science, The National University of Singapore and Charles Perkins Centre for their support and intellectual engagement. Special thanks to Yingxin Lin for the feedback on **Fig. 1**, Daniel Kim and Eugenie Lee for the active spatial omics discussion. Additional thanks to Junbin Gao for his contribution to the GTEx discussion and Boxiang Liu’s team for their insightful discussions at the Department of Pharmacy and Pharmaceutical Sciences, Faculty of Science, National University of Singapore, Singapore.

This work is supported by USYD-NUS Ignition grant to A.Z., B.B., L.Y., H.R., Y.D., Y.M., B.L. and J.Y.H.Y.; Research Training programme (RTP) Scholarship to A.Z. and S.Y.; NHMRC Investigator Grant APP2017023 to L.Y and J.Y.H.Y.; Judith and David Coffey Life Lab (CPC) to J.Y.H.Y.; Charles Perkins Centre’s Jennie Mackenzie Research Fund to Y.C. The funding source had no role in the study design; in the collection, analysis, and interpretation of data, in the writing of the manuscript, or in the decision to submit the manuscript for publication.

## Author Contributions Statement

JY conceived and led the study with design input from AZ, LY, HR and BB. AZ led the method development and implementation of the framework, with input from SY, LY, JK and JY. AZ and LY led the data assessment with input from JY, YC and JK. AZ and LY led the evaluation with input from JY, YC, EP and JK. LY and BB led the cell annotations with input from AZ. YM and HR led the nuclei segmentation dataset with input from AZ. BB led the GTEx analysis with input from JY, LY, YD, BL and AZ. EH led the imputation benchmark with input from JY, JK and AZ. All authors contributed to the writing, editing, and approval of the manuscript.

## Materials and Methods

### Datasets and preprocessing

#### Subcellular in situ spatial transcriptomics datasets from 10X Xenium

The Xenium datasets listed below were used to train GHIST+, with specific training and testing splits described under **Model Implementation**.

i. Breast cancer
  - BreastCancer1 and BreastCancer2 https://www.10xgenomics.com/products/xenium-in-situ/preview-dataset-human-breast
  - BreastCancerIDC https://www.10xgenomics.com/datasets/xenium-ffpe-human-breast-with-custom-add-on-panel-1-standard
  - BreastCancerIDC2 and BreastCancerILC https://www.10xgenomics.com/datasets/ffpe-human-breast-with-pre-designed-panel-1-standard
  - BreastCancerIDC3 https://www.10xgenomics.com/datasets/ffpe-human-breast-with-custom-add-on-panel-1-standard
ii. Colorectal cancer
  - ColorectalNormal and ColorectalADC https://www.10xgenomics.com/datasets/human-colon-preview-data-xenium-human-colon-gene-expression-panel-1-standard
  - ColorectalADC https://www.10xgenomics.com/datasets/ffpe-human-colorectal-cancer-data-with-human-immuno-oncology-profiling-panel-and-custom-add-on-1-standard
iii. Lung cancer
  - LungADC https://www.10xgenomics.com/datasets/preview-data-ffpe-human-lung-cancer-with-xenium-multimodal-cell-segmentation-1-standard
  - LungNSCLC https://www.10xgenomics.com/datasets/ffpe-human-lung-cancer-data-with-human-immuno-oncology-profiling-panel-and-custom-add-on-1-standard
  - LungADC2 https://www.10xgenomics.com/datasets/xenium-human-lung-preview-data-1-standard
iv. Pancreatic cancer
  - PancreasADC https://www.10xgenomics.com/datasets/ffpe-human-pancreas-with-xenium-multimodal-cell-segmentation-1-standard
  - PancreasADC2 https://www.10xgenomics.com/datasets/ffpe-human-ductal-adenocarcinoma-data-with-human-immuno-oncology-profiling-panel-1-standard
  - PancreasADC3 https://www.10xgenomics.com/datasets/pancreatic-cancer-with-xenium-human-multi-tissue-and-cancer-panel-1-standard

The data preprocessing protocol followed the same workflow as our previous GHIST paper ^5^ and was applied across all subcellular datasets. Briefly, we (i) parsed Xenium platform provided output folder to obtain H&E image, per-cell coordinates and gene matrices, (ii) aligned the H&E image with its corresponding coordinate frame, (iii) segmented nucleus using HoverNet pretrained on the CoNSeP dataset^39^and matched each segmented cell to its expression profile and (iv) applied the same cell annotation framework scClassify to produce aligned H&E image with its nuclei segmentations, cell gene matrix and matched nuclei coordinates for model training and evaluation. All slides above are used as input to train GHIST+. Table 1 summarizes the processed datasets used in this study, including the number of detected cells and its number of genes

#### GTEx whole slide histopathological images of breast

GTEx whole slide histopathological images of breast are retrieved from https://gtexportal.org/home/histologyPage. Nuclei segmentation was performed using HoverNet (trained on CoNSep) with pre-trained weights in original mode and nuclei type classification disabled (nr_types = 0). Whole-slide images (Aperio SVS) were tessellated into non-overlapping 2000 × 2000 pixel patches, with right-edge, bottom-edge, and corner patches added to ensure full coverage. No stain normalisation or color augmentation was applied.

Each 2000 × 2000 patch was processed using a sliding-window strategy with 270 × 270 pixel input tiles (native resolution in original mode) and an 80-pixel stride, yielding 190-pixel overlaps.

Reflection padding was applied at patch boundaries. Each tile produced an 80 × 80 pixel segmentation output. Inference used a batch size of 128 tiles, with 8 data loading workers and 32 post-processing workers per GPU. The model generated instance-level segmentation maps with unique integer IDs per nucleus.

Overlapping tile predictions were first merged within each patch. Patch-level maps were then combined into a whole-slide segmentation by reassigning globally unique IDs. Nuclei split across 2000 × 2000 patch boundaries were detected using outer boundary extraction (skimage.segmentation.find_boundaries, mode = ‘outer’); nuclei within 10 pixels of patch edges were examined for cross-patch overlap and merged by retaining the lower ID.

#### scRNA-seq reference datasets used in cell-type classification

For each dataset, we inherited the processing from the GHIST paper and used scClassify^40^ (version 1.22.0) to annotate the cell types of each cell. This method uses reference data with known cell types for classification.

##### Breast cancer reference

The reference dataset was based on a publicly available breast cancer dataset from 10x Genomics. The dataset was processed following the same pipeline described in our previous publication^41^. Original dataset download link: https://www.10xgenomics.com/products/xenium-in-situ/preview-dataset-human-breast/

##### Colon cancer reference

The reference dataset was obtained from the study Integrative single-cell analysis of human colorectal cancer reveals patient stratification with distinct immune evasion mechanisms^42^. Only tumour tissue and normal adjacent tissue were retained, and the cell types ILC, Proliferating T, and Proliferating Myeloids were excluded. Original dataset download link: https://doi.org/10.6084/m9.figshare.25323397

##### Lung cancer reference

The reference dataset was constructed from the single-cell lung cancer atlas (LuCA) core atlas, selecting only adenocarcinoma samples to build reference cell-type profiles^43^. Only lung adenocarcinoma samples from primary tumour tissues were retained. Original dataset download link: https://cellxgene.cziscience.com/collections/edb893ee-4066-4128-9aec-5eb2b03f8287

##### Pancreatic cancer reference

The reference dataset was derived from the human pancreatic cancer single-cell atlas dataset^44^. Only cells annotated as primary tumour were retained, and clusters labelled as cycling were excluded. Original dataset download link: https://zenodo.org/records/14199536

##### Normal breast tissue reference

The reference dataset was obtained from the Human Breast Cell Atlas^45^. The dataset is publicly available at: https://cellxgene.cziscience.com/collections/48259aa8-f168-4bf5-b797-af8e88da6637

#### Overview of GHIST+

Current H&E-to-SGE advances leave behind a central challenge of core-to-WSI reconstruction: using sparsely profiled tissue cores to infer cell-level molecular states across WSIs. This requires a model that uses local cellular neighbourhood context to predict cell-level molecular states beyond profiled cores, learns recurring tissue states to connect sparse cores with unprofiled WSI regions, and handles heterogeneous gene-panel coverage by imputing genes unobserved from a given panel. These requirements motivate a generalisable framework for tissue-wide molecular reconstruction from sparse and heterogeneous spatial measurements.

GHIST+ is a generalisable tissue wide molecular reconstruction framework capable of training on multiple H&E stained histology images for per-cell expression prediction across slides, tissue types and gene panels. GHIST+ combines three modules: (i) an Edge-conditioned Residual Mixer (ECRM) that propagates local cellular neighbourhood context at true cell resolution, (ii) vector-quantised (VQ) prototyping that learns recurring tissue states to connect sparse cores with unprofiled WSI regions, (iii) a Mask-Aware Gene Conditioned Imputer (MAGCI) for imputing genes unobserved from a given panel and unifying heterogenous panels across slides (**Fig. 1**).

Complete details on the model implementation, including parameter selection and training configuration, will be released in our GitHub repository https://github.com/SydneyBioX/GHIST_plus.

GHIST+ initially extracts tile level and cell-level morphology features from each 224 ×224 histology patch using the UNI2h pathology foundation model. For each tile *b*, the encoder produces two feature maps *hd*1_*b*_(*u*) and *h*1_*b*_(*u*) over pixel locations *u*. We summarize tile level features by mean pooling over the tile and cell-level morphology within each nucleus mask *M*_*i*_as follows:

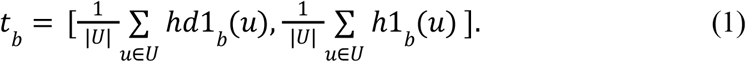

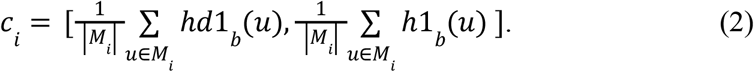

We then calculate the final per-cell morphology vector *𝒱*_*i*_ by concatenating cell summary *c*_*i*_ with tile summary *t*_*b*_.

### ECRM

ECRM is a graph-based module that refines each cell representation using information from local tissue neighbourhoods through three steps: (i) selection of relevant neighbours (ii) building local cell context (iii) using local context for expression reconstruction (**Fig. 1b**).

#### (i) Selection of relevant neighbours

Starting from the morphology-derived cell state 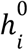, GHIST+ builds a local graph around each cell *i*. For each neighbouring cell *j* ∈ *N*(*i*), the edge vector *g*_*ij*_ describes how the neighbour relates to the target cell, including spatial distance, relative position, morphology similarity and cell-type agreement. ECRM converts this edge vector into a neighbour weight 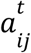, and uses these weights to form a context summary 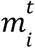 from nearby cell states below:

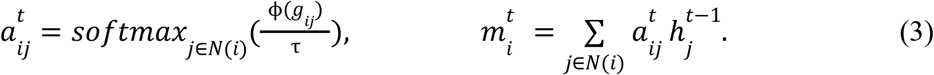

where ϕ() denotes a learnable edge scoring function that maps *g*_*ij*_ to an unnormalised neighbor score and τ is a temperature parameter controlling neighbour weighting.

#### (ii) Building local cell context

ECRM then updates each cell by mixing its own state with the neighbour summary. A learned gate controls how much neighbourhood information is added, allowing the model to use local tissue context while retaining the cell’s own morphology signal.

#### (iii) Using local context for expression reconstruction

The ECRM refined representation is then used to estimate reference-state weights for each cell. Given reference expression vectors *R*_*r*_, GHIST+ constructs a reference-guided baseline expression 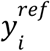 as follows:

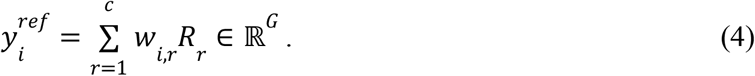

This baseline captures expression from inferred cell state. GHIST+ then predicts pre-imputation expression by adding two learned corrections: a composition-conditioned correction 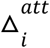 and a graph refined morphology correction 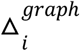. The pre-imputation prediction is then calculated as follows:

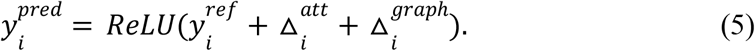

### VQ

VQ is a module in GHIST+ that groups tiles with similar morphology into a small set of learned tissue prototypes. These prototypes are used in two ways: (i) provide tile-level context for expression prediction and (ii) select representative TMA regions for broadcasting (**Fig. 1b**).

#### (i) Provision of tile-level context

##### a Learning repeated tissue patterns

Each tile summary *t*_*b*_ from equation (1) is passed through a learned projection layer, resulting in tile vector *z*_*b*_. VQ then matches *z*_*b*_ to the closest learned prototype, producing quantised tile representation 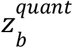 and a prototype code *q*_*b*_. The VQ loss learns these prototypes and keeps each tile close to its assigned prototype. *sg* here denotes stop gradient. The equation is as of follows:

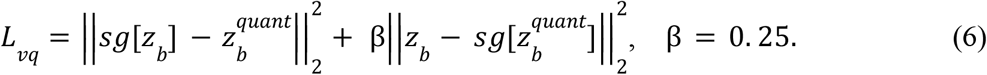

This step allows GHIST+ to represent repeated tissue patterns using shared prototypes.

##### b Integrating tile context

The quantised tile representation 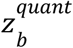 is used as tile context for expression prediction. It is passed to cells within the same tile, allowing each cell prediction to use morphology and tissue patterns around it.

#### (ii) Selection of representative TMA regions

The same prototype codes are reused to select representative TMA cores. For each candidate region centered at *c*, GHIST+ calculates the proportion of each prototype code *k* among quality controlled tiles *W*_*qc*_ (*c*) where,

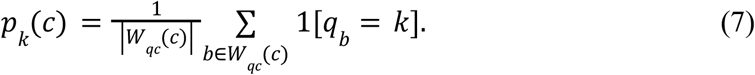

This produces prototype profile *p*(*c*) which summarizes the tissue patterns inside the candidate region. GHIST+ then scores each region by comparing its prototype profile with target profile *t*, while also accounting for tissue coverage and region size according to:

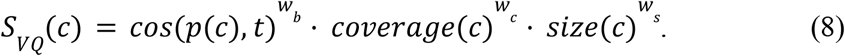

The highest scoring candidate ROI is selected as the representative region for broadcasting. Candidate ROIs can be further reranked using additional biological criteria: cell type composition and cell density.

### MAGCI

MAGCI is a module in GHIST+ that unions and imputes unobserved genes. During training, a subset of observed genes is randomly hidden and the model learns to reconstruct them from retained genes with the aid of morphologically derived cell context from pre imputation prediction 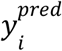 (**Fig. 1b**). As a result, the imputer learns to recover unobserved signals under panel mismatch.

The final imputed expression is obtained by adding the imputed residual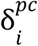 to the reference baseline 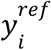 in equation (4). The final equation is as of follows:

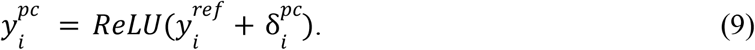

The overall per-cell imputed expression output is defined as 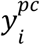 after panel completion.

### Model Optimisation

We optimized GHIST+ with a weighted multitask loss over mini-batches of 64 cells. The objective adopted a multitask training structure with adaptation to support imputation, prototyping and pan-cancer training. The total losses are grouped into four functional loss categories: expression supervision, cell type supervision, composition regularisation and auxiliary supervision. The equation is as of follows:

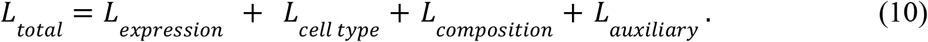

The expression loss supervises observed gene expression using a zero-aware error and masked correlation loss, with additional expression supervision for immune and invasive cell types. The cell-type supervises two classification heads: one using histology features and one using predicted expression. Their embeddings and logits are aligned so both heads predict consistent cell identities. The composition loss matches predicted cell-type proportions at the tile and mini-batch levels to the observed proportions.. The auxiliary term includes hidden gene imputation for panel completion and VQ prototype loss in equation (6). Loss ablations compared to GHIST is shown in **Supplementary Fig. S11**.

### Model Implementation

During training, GHIST+ is instantiated with a frozen UNI2h backbone that extracts histology features from each image patch. Per-cell embeddings were obtained through mean-pooling encoder features within each nuclei mask and concatenating them with a globally pooled tile representation. Neighbourhood context is incorporated with a two step ECRM operating with a centroid-based kNN neighbourhood cell graph (*k* = 12 within patch) augmented with cross patch edges (*k* = 8 across patch). Pooled tile features are discretized into patch prototypes (64 codes) which provide contextual cues during expression prediction. Finally, all samples were mapped into a shared union gene space with a binary gene mask and slide identity that allow GHIST+ to use the correct slide specific reference matrix for each batch.

For all experiments, five-fold cross validation was performed within each slide such that 80% of each Xenium slide is used for training and 20% for validation in each run, while across-slide experiments held out one entire slide as an unseen external test set. Histology images were tiled into 224 ×224 patches with no overlap during training and a 30 pixel overlap during validation and external inference. Patches were filtered where the minimum number of nuclei within a patch must be greater or equals to one. H&E histology preprocessing includes Macenko stain normalisation ^46^ against a reference H&E stain (See ablation study in **Supplementary Fig. S12**) and per-channel RGB standardisation. Additional on-the-fly augmentation during training include flipping (horizontal/vertical), rotations (90/180/270 degrees) and HED-based colour perturbation ^47^. Training utilized learning rate of 10^−3^, first-moment estimate of 0. 9, second-moment estimate of 0. 999 and weight decay of 10^−4^.

GHIST+ was trained for 30 epochs with a batch size of 64. We ran GHIST+ on a 48GB NVIDIA GeForce RTX A6000 GPU, Intel Xeon CPU with 16 cores, 32 threads and 502 GB RAM. Training was separated into three categories: (i) Breast slide used in **Fig. 3a** and **b** (ii) Breast slides used in **Fig. 2, Fig. 3c**-**f** and **Fig. 4** and (iii) Pan-cancer slides used in **Fig. 5**.

i. Breast slide
  - Training completed within 2 hours using Xenium BreastCancer2 slide.
  - Training area includes 80% of the slide region.
  - Inference area includes 20% of the slide region.
ii. Breast slides
  - Training completed within 72 hours using all Xenium breast slides (five slides for training one slide for testing).
  - Training slides include BreastCancer1, BreastCancerIDC, BreastCancerIDC2, BreastCancerILC and BreastCancerIDC3.
  - Xenium BreastCancer2 was used as an independent inference slide.
iii. Pan-cancer slides
  - Training completed within 120 hours using Xenium Pan-cancer slides (breast/pancreas/colorectal/lung).
  - Training slides include all slides except PancreasADC3.
  - Inference slides include PancreasADC3.

### Comparison methods

We compared GHIST+ with state of the art methods GHIST and SpatialEx. H&Enium was not compared due to a lack of code release at the time of writing.

### Evaluation study 1: Method comparisons at single-cell resolution

In our study, we ranked genes according to Giotto for SVGs and Scanpy for HVGs. We evaluated GHIST+ with two types of metric categories: gene expression accuracy and biology recovery.

### Gene ranking and expression prediction

To rank SVGs, we followed GHIST and scored spatially variable genes using Giotto calculation where we let *g* represent a gene, 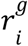 represent the ranked expression of a gene *g* in a cell *i*and 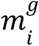 be the mean ranked expression of gene *g* across k nearest neighbours of cell *i*. We scored each gene *g* with a *score*_*g*_ and ranked them in descending order, extracting the top 20 and 50 as SVGs:

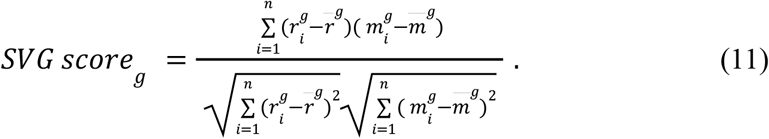

To rank HVGs, we scored highly variable genes using Scanpy from Python where we score each gene *g* with bin-normalized dispersion for ranking. δ_*g*_ here represents the log dispersion of gene *g* and *B*(*g*) is the mean-expression bin containing gene *g*. We scored each gene *g* with *score*_*g*_ and ranked them in descending order, extracting the top 20 and 50 HVGs:

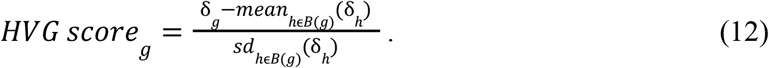

In our evaluation pipeline, PCC is calculated between the ground truth genes and predicted genes across all cells. We define *x*_*ig*_ as the ground truth expression of gene *g* in cell *i*and 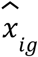 as the predicted expression of gene *g* in cell *i*. We then calculate PCC per gene *g* across the total number of cells:

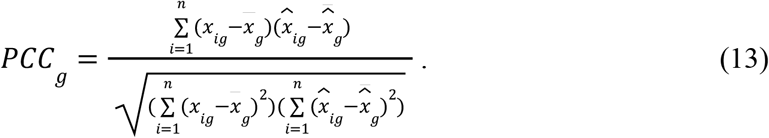

Spearman’s correlation mean is calculated for each gene as the Pearson correlation between the ranked ground truth and ranked predicted expression values across cells. Here, *R*_*ig*_ and 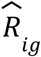 represents the ranked ground truth and predicted expression values of gene *g* in cell *i*. The equation is as of follows:

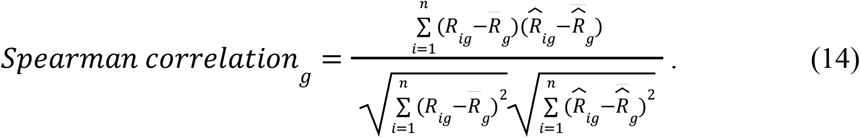

Root Mean Squared Error is calculated per gene across overlapping cells after *log* normalisation of ground truth expression *x*_*ig*_ and predicted expression 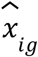 of gene *g* per-cell *i. l i*symbolize the ground truth size of *i*and *n* represents the number of cells. The equation is as of follows:

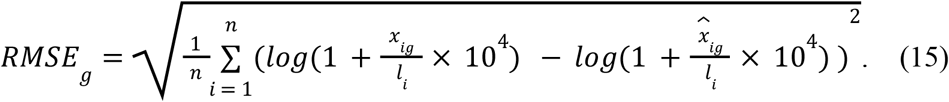

### Biological and spatial recovery

SSIM score is calculated for per gene *g* where *G*_*g*_is defined as the ground truth spatial expression map and *P*_*g*_ is the predicted spatial expression map. Both metrics were generated by spatial binning of cell expression, Gaussian smoothing and cropping to the tissue area. µ, σ^2^ and 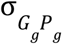 denote the corresponding mean, variance and covariance. The equation is as of follows:

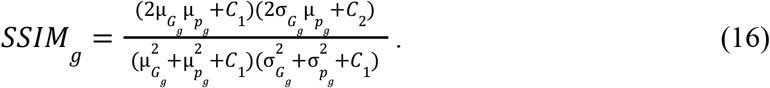

Correlation Matrix Distance is used to assess preservation of gene structure between predicted expression *R*_*predicted*_ and ground truth *R*_*true*_ :

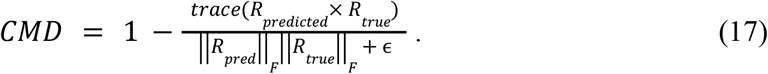

TISSUE score is computed for each cell to gene pair as the signed prediction error normalized by a variability term ^17^. *X*_*ij*_ and 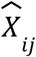 denote the observed and predicted expression of gene *j* which calculates *U* that represents local prediction variability across neighbourhoods *N*_*ij*_ weighted by *W*_*ik*_. The equation is as of follows:

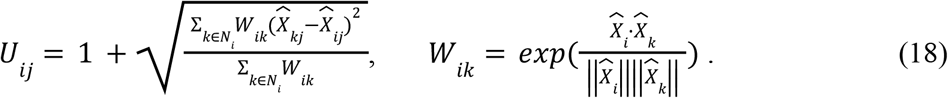

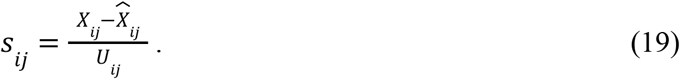

For predicted expression, we calculate its cell type marker recovery using pyUCell v0.5.0 ^48^. For each cell, pyUCell rank-based signature scores were computed from curated marker sets for B, epithelial, macrophage, malignant, myeloid, plasma, stromal and T cell populations, using *max*_*rank* equal to the number of shared genes, *missing*_*genes* = *“;skip”*, and a chunk size of 500. The we defined unassigned cell type score as 1 − *maximum score* across all biological cell-type signatures.

AUROC is computed for each cell type in one versus the rest manner using rank-based formulation. We let *n*^+^ denote positive samples, *n*^−^ denote the negative samples and *R*^+^ denote the sum of the ranks of the predicted scores assigned to positive samples after ranking all scores. The AUROC equation is as follows:

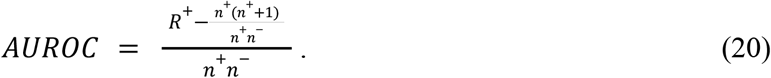

Recall is computed to summarize cell type recovery where *C* defines the number of cell type classes, *N*_*cc*_ represents the number of cells that have true and predicted label as *c* whereas *N*_*cj*_ represents the number of cells that have true cell type label as *c* but predicted cell type as *j*. The overall equation is as follows:

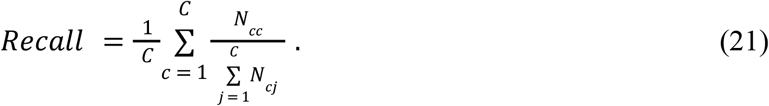

For spatial evaluation, each cell type was considered independently. Cells were ranked by their pyUCell score for a given class, and the top *n* cells were selected, where *n* matched the number of ground-truth cells of that class. Predicted and ground-truth positive cells were converted into smoothed spatial density maps using 150 μm bins and Gaussian smoothing with sigma 1.2 bins. Spatial agreement was quantified by the intersection-over-union between ground truth spatial bin 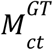 and predicted spatial bins 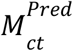 of each cell type *ct* as follows:

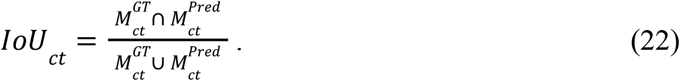

### Evaluation study 2: Imputation analysis

For the masked imputation benchmark, we selected the top 10 Giotto-ranked SVGs from each slide and treated them as unmeasured genes by masking them from the observed-gene input and training loss. GHIST+ was then evaluated by comparing the reconstructed expression of these held-out genes with their original observed expression. Evaluation of imputation performance adheres to metrics defined under **Evaluation study 1**. Additional method description is under **Overview of GHIST+**.

### Evaluation study 3: Pathway enrichment analysis

#### Spatial pathway enrichment analysis

Pathway activity at the single-cell-level was estimated using single-sample gene set enrichment analysis (ssGSEA). Here, each cell is a single sample. Hallmark gene sets were obtained from the MSigDB H collection using the msigdbr R package. ssGSEA scores were computed with the GSVA package using log-normalized gene expression values. To investigate local pathway patterns, several regions of interest were manually defined based on spatial coordinates: a top-left region (x = 8100-11500, y = 8200-11000), a bottom-left region (x = 6300-9700, y = 15500-18200), and a bottom-middle region (x = 14100-17500, y = 16300-19000). Cells falling within each region were identified, and the median ssGSEA score of each pathway was calculated for that region. For each region, pathways were ranked by their median enrichment score and the top pathways were visualised using bar plots.

### Whole slide spatial pathway enrichment maps

Whole slide spatial pathway enrichment maps were generated by aggregating single-cells into spatial grids and computing pathway activity within each grid. Briefly, cells were grouped into square spatial bins of 500 pixels (∼106 μm) based on their spatial coordinates. For each grid region, gene set enrichment scores were calculated using the GSVA method on log-normalized gene expression values. Hallmark gene sets from the MSigDB H collection were used as pathway references. Regions containing fewer than 10 cells were excluded to ensure robust estimation of pathway activity. For comparison between ground-truth and predicted datasets, Pearson and Spearman correlations were computed across regions.

### Statistical analysis on GTEx breast tissues data

#### Cell type annotation

Cell type annotation for GTEx whole slide histopathological images of breast was performed using a two-stage hierarchical classification strategy implemented with scClassify ^40^. In the first stage, a broad classifier was trained to distinguish three major cell types (epithelial, stromal and immune cells) using the Human Breast Cell Atlas (HBCA) reference dataset ^45^. Cells annotated as doublets or stripped nuclei were excluded from the training set. To balance class representation, we randomly subsampled 5,000 cells per class (15,000 cells in total). Model training was restricted to genes overlapping with those available in the predicted GTEx spatial transcriptomics data.

In the second stage, we refined immune cell annotations by training a classifier on immune cells only. Specifically, cells labelled as immune in the first-stage reference were subsetted, and a secondary model was trained to distinguish B cell and T cell populations. Given the limited gene overlap (∼1,000 genes) between the reference and GTEx data, we only focused on B cells and T cells.

For prediction, the hierarchical model was applied sequentially. All cells were first assigned to one of the three major compartments using the primary classifier. Cells predicted as immune were subsequently passed to the second-stage classifier to obtain refined immune subtype labels (B or T cells), while non-immune cells retained their original epithelial or stromal annotations.

#### Spatial niche detection using BANKSY on normal breast tissue

Spatial niches in GTEx normal breast tissue were identified using the BANKSY framework. Briefly, dimensionality reduction was performed using principal component analysis (PCA), retaining the top 30 principal components. Clustering was subsequently performed on the BANKSY-transformed representation, and five spatial niches were identified (n = 5 clusters). Default settings were used for all other parameters unless otherwise specified.

Further niche detection was performed within each compartment of the GTEx normal breast tissue using the BANKSY framework in a similar way using the following setting. Cells were first subset to the epithelial compartment. BANKSY neighbourhood features were computed separately within each sample using computeBanksy with assay_name = “logcounts” and k_geom = 10. Per-sample BANKSY objects were then combined, followed by PCA using runBanksyPCA with lambda = 0.8, npcs = 30, and group = “sample_id”. Clustering was performed using clusterBanksy with lambda = 0.8, npcs = 30, algo = “kmeans”, and kmeans.centers = 6, with set.seed(123). The resulting epithelial subniches were used for marker ranking, cell-type composition summaries, and age-associated ECM-remodelling analysis.

### Supervised histology-guided compartment

A supervised histology-guided classifier ^18^ using code from GitHub (https://github.com/cancerbioinformatics/OASIS) was used to assign tissue regions into canonical epithelial, stromal, and adipose compartments based on histological morphology.

#### Age-associated analysis

### To assess age-associated changes in cellular composition and microenvironmental programmes, we modelled age as a continuous variable across GTEx samples using linear regression models in R. The following phenotypes were evaluated

(a)Cellular composition: For each sample, cell-type proportions were calculated as the fraction of cells assigned to each cell type relative to the total number of cells within the sample or within each spatial niche. Immune-cell proportions were computed using predicted B-cell and T-cell annotations derived from GHIST+ cell-type predictions.

(b)ECM-remodelling score: We calculated a targeted ECM remodelling score as the mean log-transformed expression of 11 curated ECM-associated and stromal-remodelling genes. These were *DCN, LUM, COL1A1, COL5A2, THBS2, MFAP5, POSTN, MMP2, MMP3, PDGFRA*, and *PDGFRB*. This averaging strategy follows previous ECM signature scoring approaches ^49,50^.

