## Supplementary material for "Generalisable tissue-wide molecular reconstruction from histology": All supplementary figures

#### Supplementary Tables

**Supplementary Table 1.** Top 10 DE genes for each three histology-defined tissue compartments.

| Compartment | Rank | Gene | LogFC (base 2) | Adjusted P-value | In compartment (mean) | Not in compartment (mean) |
|---|---|---|---|---|---|---|
| **Epithelial** | 1 | WFDC2 | 0.76 | $1.92 \times 10^{-34}$ | 1.05 | 0.30 |
| | 2 | TACSTD2 | 0.68 | $9.65 \times 10^{-24}$ | 4.29 | 3.61 |
| | 3 | KRT7 | 0.63 | $2.30 \times 10^{-25}$ | 3.96 | 3.33 |
| | 4 | AZGP1 | 0.57 | $9.17 \times 10^{-32}$ | 1.87 | 1.30 |
| | 5 | CD37 | 0.50 | $6.05 \times 10^{-43}$ | 0.85 | 0.35 |
| | 6 | CLDN4 | 0.48 | $7.15 \times 10^{-24}$ | 2.96 | 2.48 |
| | 7 | CCL28 | 0.44 | $1.82 \times 10^{-38}$ | 0.96 | 0.52 |
| | 8 | KRT8 | 0.40 | $7.79 \times 10^{-22}$ | 3.51 | 3.11 |
| | 9 | KRT15 | 0.37 | $2.89 \times 10^{-30}$ | 1.08 | 0.71 |
| | 10 | KRT14 | 0.36 | $5.61 \times 10^{-26}$ | 2.14 | 1.77 |
| **Stromal** | 1 | IGFBP7 | 0.82 | $3.47 \times 10^{-29}$ | 6.28 | 5.45 |
| | 2 | SPARCL1 | 0.64 | $1.63 \times 10^{-29}$ | 4.60 | 3.95 |
| | 3 | SERPINE1 | 0.62 | $1.66 \times 10^{-29}$ | 3.44 | 2.81 |
| | 4 | ANGPT2 | 0.62 | $9.13 \times 10^{-30}$ | 3.80 | 3.18 |
| | 5 | IFI27 | 0.61 | $1.27 \times 10^{-29}$ | 3.45 | 2.84 |
| | 6 | TCF4 | 0.56 | $3.81 \times 10^{-29}$ | 3.98 | 3.42 |
| | 7 | CLDN5 | 0.54 | $6.93 \times 10^{-30}$ | 2.70 | 2.16 |
| | 8 | GNG11 | 0.53 | $8.10 \times 10^{-30}$ | 3.37 | 2.84 |
| | 9 | STC1 | 0.52 | $2.16 \times 10^{-29}$ | 3.84 | 3.32 |
| | 10 | CAV1 | 0.50 | $5.30 \times 10^{-29}$ | 4.49 | 3.99 |
| **Adipose-associated** | 1 | IGFBP7 | 0.45 | $1.49 \times 10^{-12}$ | 6.03 | 5.58 |
| | 2 | SPARCL1 | 0.32 | $6.21 \times 10^{-11}$ | 4.38 | 4.06 |
| | 3 | CD74 | 0.31 | $3.73 \times 10^{-20}$ | 6.41 | 6.10 |
| | 4 | SERPINE1 | 0.29 | $4.47 \times 10^{-10}$ | 3.22 | 2.92 |
| | 5 | TCF4 | 0.28 | $3.42 \times 10^{-11}$ | 3.79 | 3.51 |
| | 6 | IFI27 | 0.28 | $8.43 \times 10^{-10}$ | 3.22 | 2.95 |
| | 7 | CAV1 | 0.27 | $5.48 \times 10^{-12}$ | 4.33 | 4.07 |
| | 8 | ANGPT2 | 0.27 | $4.17 \times 10^{-18}$ | 3.56 | 3.30 |
| | 9 | CDKN1A | 0.27 | $5.99 \times 10^{-9}$ | 3.19 | 2.93 |
| | 10 | DCN | 0.25 | $1.98 \times 10^{-27}$ | 2.69 | 2.44 |

**Supplementary Table 2.** Age-associated epithelial microniches identified by secondary niche analysis within histology-defined epithelial compartment

| Microniche | Name | Marker genes | Beta | P-value |
|---|---|---|---|---|
| 1 | Vascular Stromal | IGFBP7,SPARCL1,ANGPT2,SERPINE1,IFI27 | 0.0008 | 0.0825 |
| 2 | Luminal Epithelial | TACSTD2,KRT7,WFDC2,MGP,CLDN4 | 0.0033 | 0.0114 |
| 3 | Immune | CD37,CXCR4,PTPRC,CD69,IL7R | 0.0012 | 0.3797 |
| 4 | ECM remodelling | IGFBP7,SERPINE1,SPARCL1,IFI27,TCF4 | 0.0019 | 0.0156 |
| 5 | Mixed immune | CXCR4,CD37,PTPRC,CD69,LAPTM5 | 0.0025 | 0.0614 |
| 6 | Luminal Epithelial | TACSTD2,KRT7,MGP,WFDC2,CLDN4 | 0.0029 | 0.0266 |

### Supplementary Figures

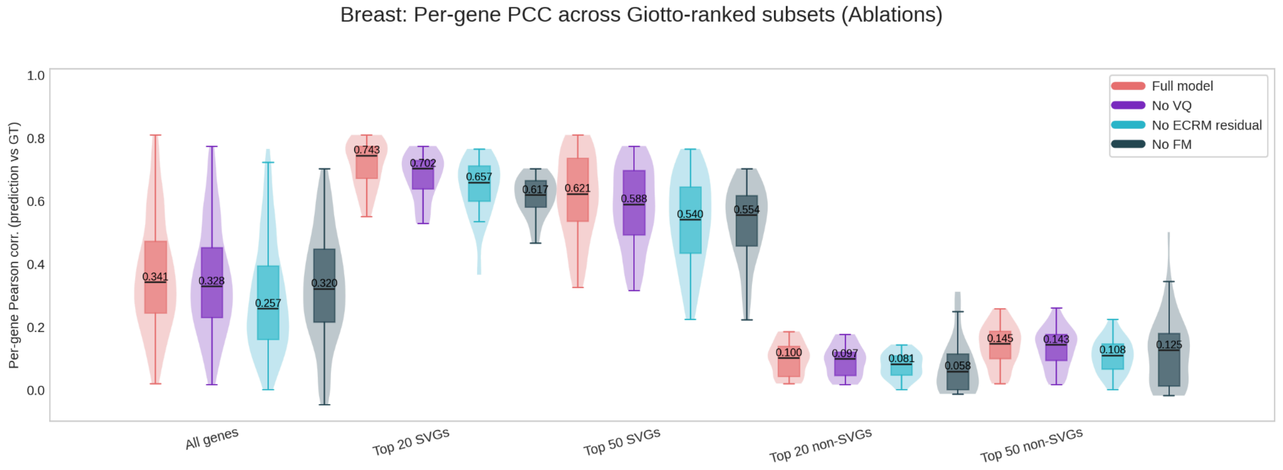


**Supplementary Figure S1: Within slide architectural ablations.**
Violin boxplot showing GHIST+ comparing variants without ECRM, VQ and foundation model trained weights. Metrics were measured across all genes, top 20 and 50 SVGs and non-SVGs. Full model GHIST+ showed the highest performance followed by an observation of decreasing performance as modules are removed.

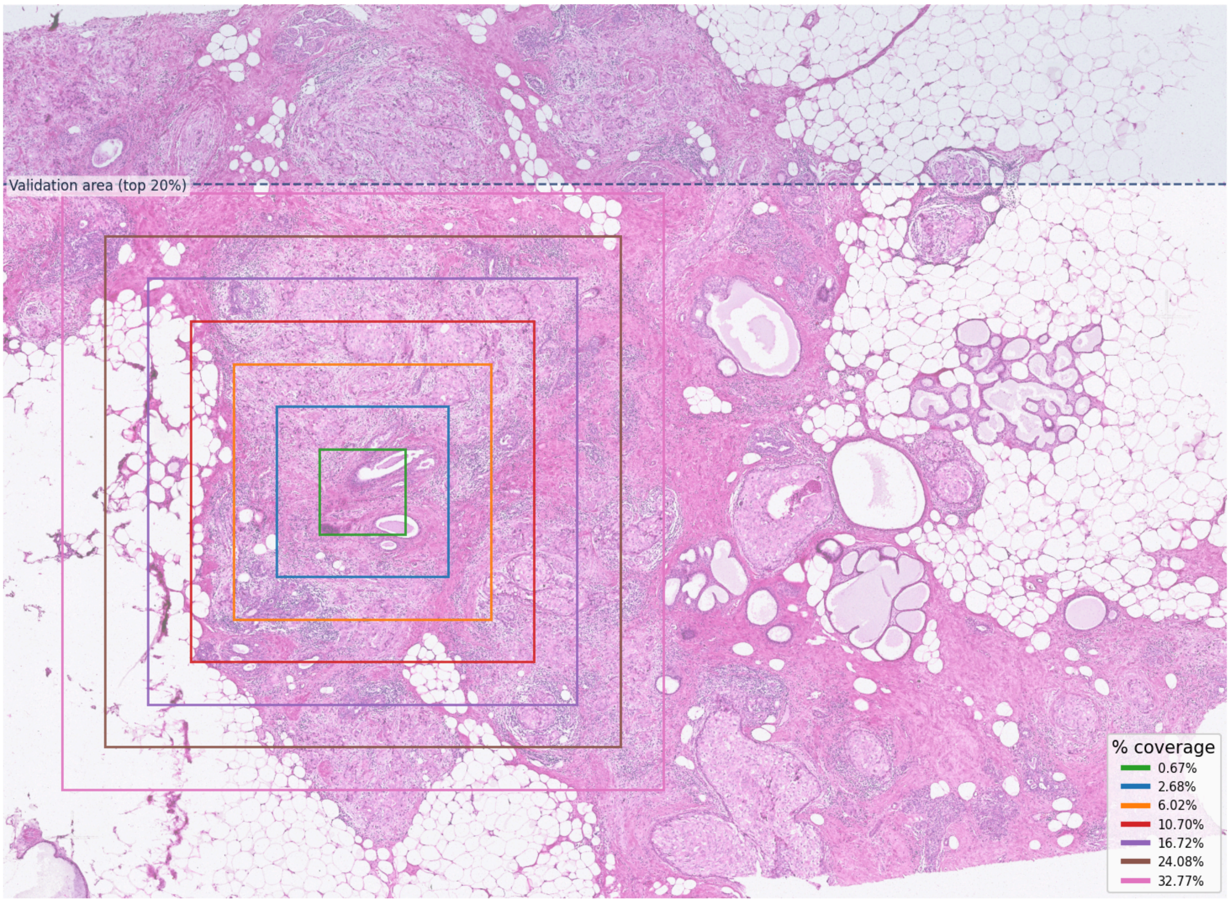


**Supplementary Figure S2: Spatial layout of the ROI saturation test.**. Experimental visualisation of a VQ selected representative ROI. The ROI is progressively expanded to simulate increasing TMA coverage, while gene prediction is evaluated in a separate held out region. This panel visualises the broadcasting design in **Fig. 4c** where limited profiled tissue is tested for its ability to support zero-shot molecular reconstruction in unprofiled tissue.

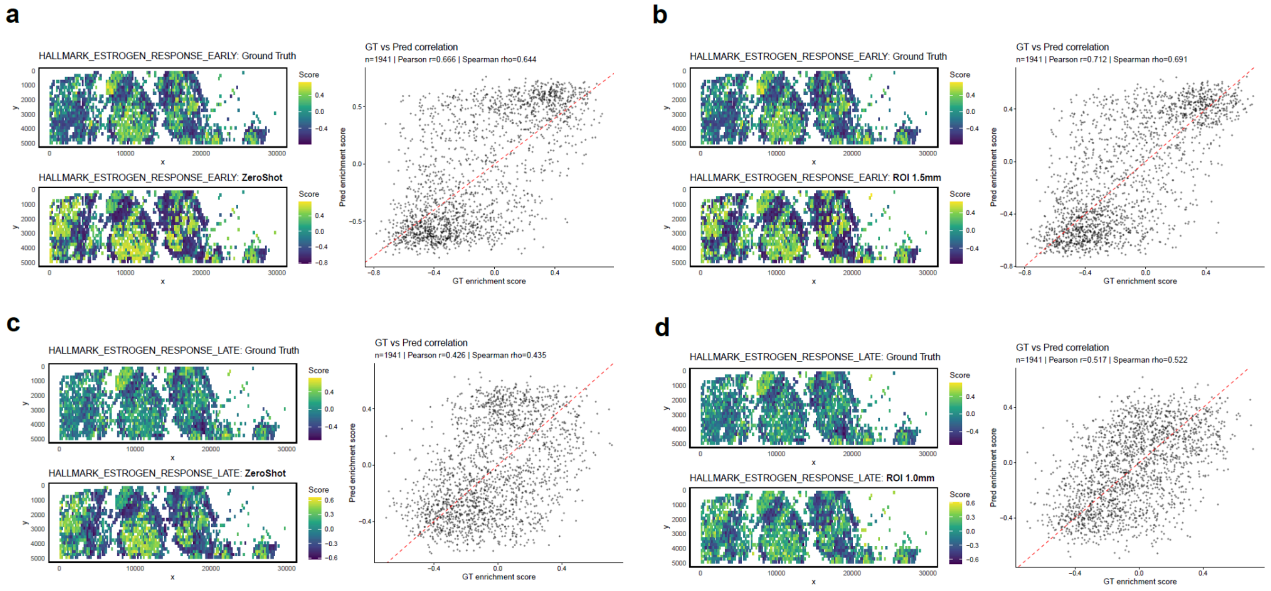


**Supplementary Figure S3: ROI-informed modelling improves recovery of spatial transcriptomic signals**. (a,c) Zero-shot inference results without TMA transcriptomic information for HALLMARK_ESTROGEN_RESPONSE_EARLY and HALLMARK_ESTROGEN_RESPONSE_LATE, respectively. (b,d) ROI-informed prediction results incorporating TMA transcriptomic context (1.5 mm ROI for HALLMARK_ESTROGEN_RESPONSE_EARLY and 1.0 mm ROI for HALLMARK_ESTROGEN_RESPONSE_LATE). For each panel, the left top shows the ground-truth pathway enrichment map derived from measured transcriptomics, the left bottom shows the enrichment map derived from predicted transcriptomics, and the right shows the correlation between ground-truth and predicted enrichment scores across spatial locations.

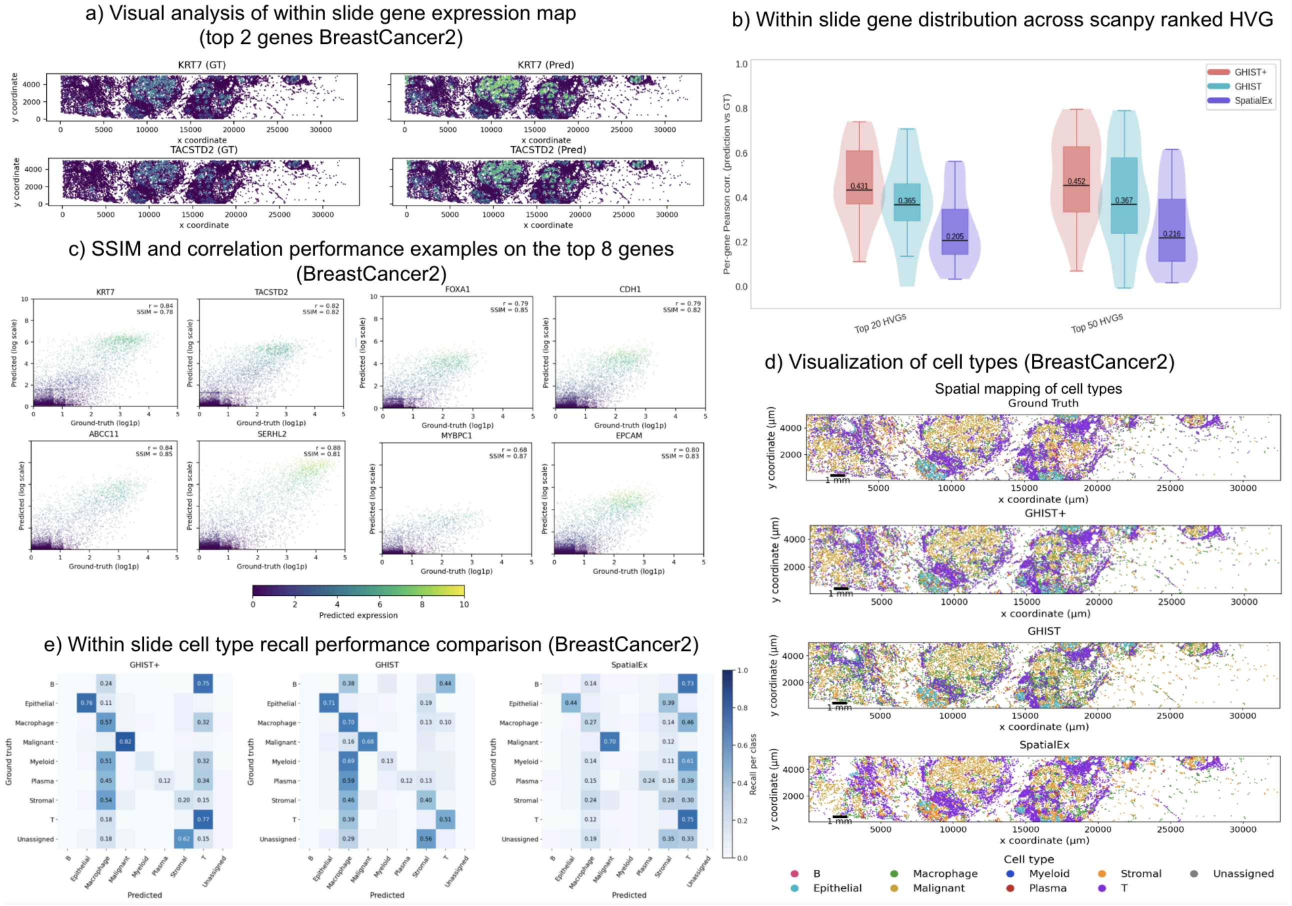


**Supplementary Figure S4: Additional within-slide evaluation of GHIST on Xenium BreastCancer2 slide.**

(a) Visual spatial map analysis of the top two genes predicted by GHIST+. (b) Violin boxplot comparison between GHIST+ with state-of-the-art subcellular gene expression prediction methods within slides using Scanpy ranked HVGs measured using per gene PCC. (c) Per-cell scatter plots for the top eight genes in BreastCancer2 compared with SSIM and r value per gene. (d) Spatial maps of cell type assignments in BreastCancer2, comparing ground truth against predictions from GHIST+ with GHIST and SpatialEx. (e) Within slide cell type recall matrices for GHIST+, GHIST and SpatialEx in BreastCancer2.

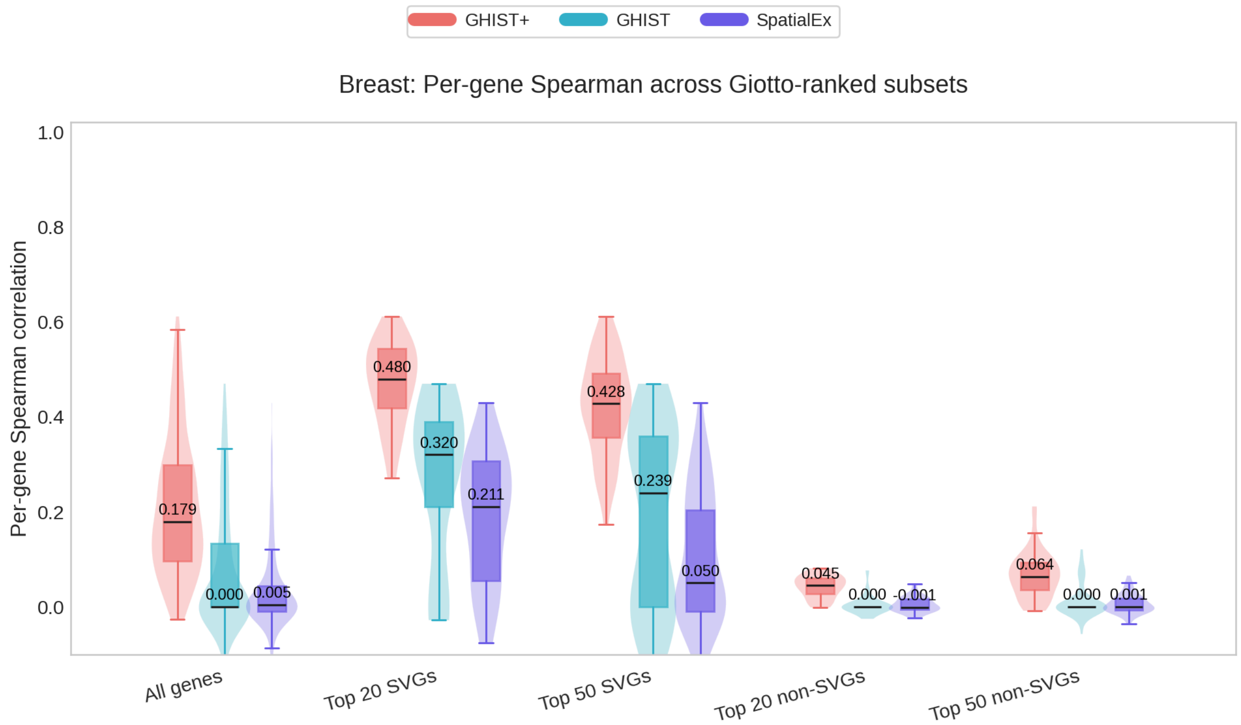


**Supplementary Figure S5. Across slide comparison of per gene Spearman correlation across Giotto-ranked gene subsets.**

Violin boxplots show per gene Spearman correlation on held out BreastCancer2 slide on all genes and top 20 and 50 SVGs and non-SVGs.

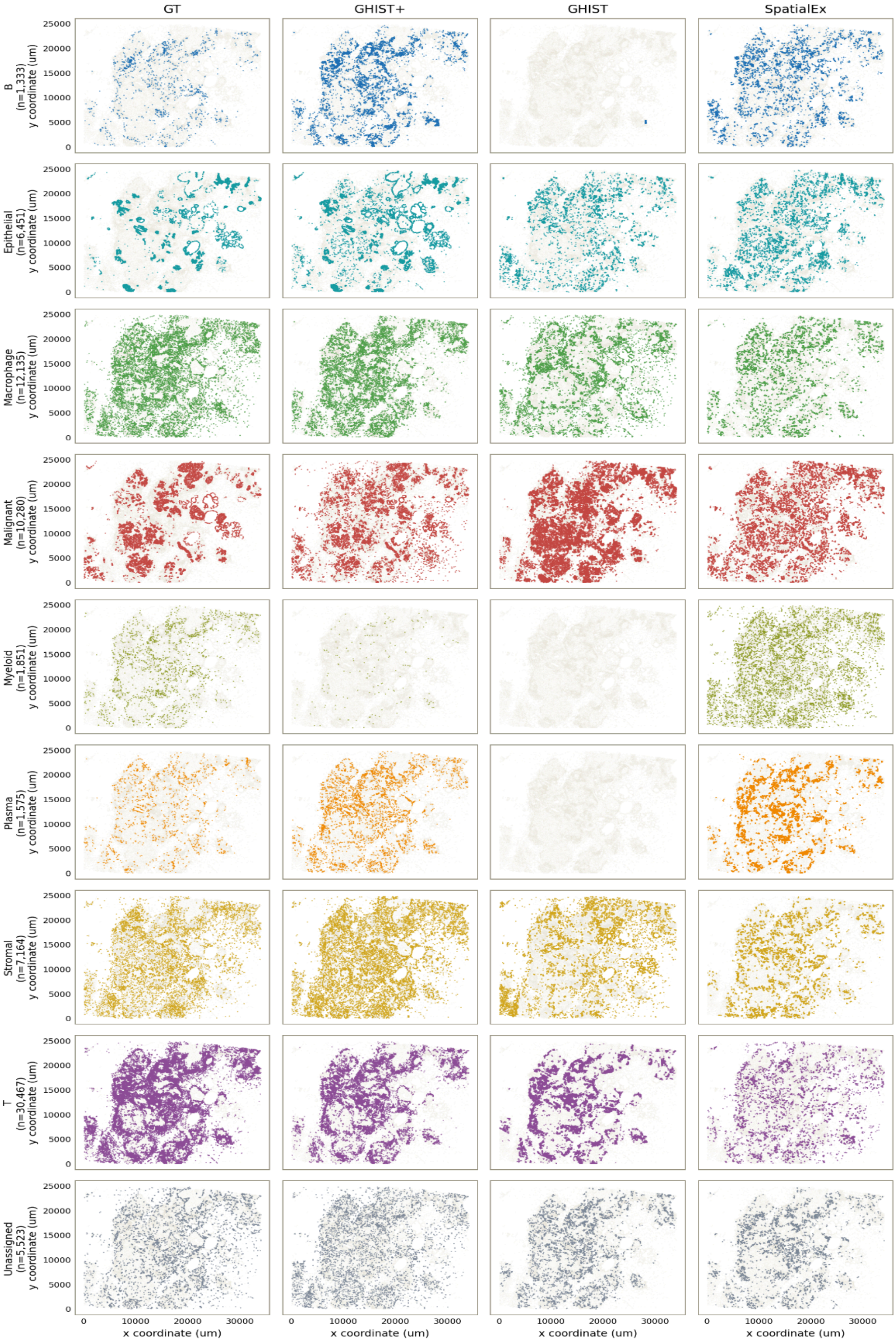


**Supplementary Figure S6: Spatial visualisation of per cell type spatial map reconstruction across GHIST+, GHIST and SpatialEx**.

Per cell type spatial maps are visually compared to the ground truth held out Xenium BreastCancer2 slide. Cell types include T cells, B cells, Epithelial cells, Plasma cells, Stromal cells, Malignant cells, Myeloid cells, Macrophages and unassigned cells. Visual comparison focuses on whether each method correctly recovers the location, shape and boundaries of cell types

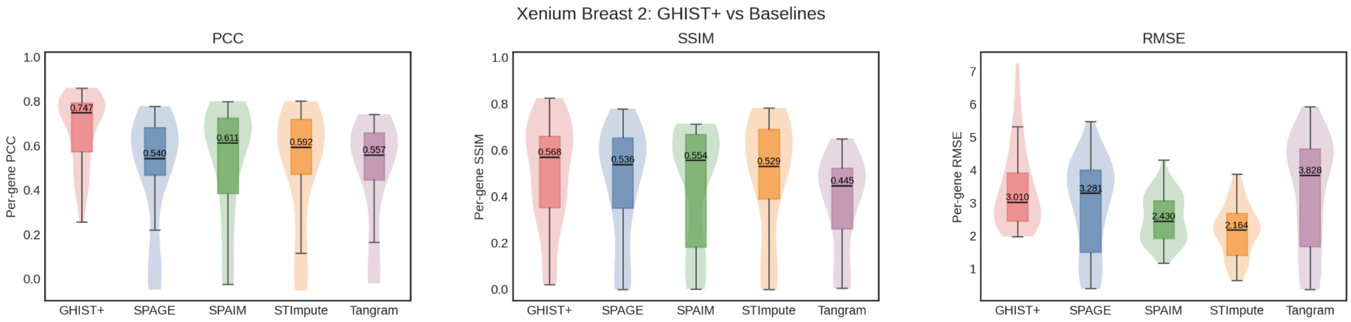


**Supplementary Figure S7: Additional method comparison with imputation algorithms.** Violin boxplots comparing performance of GHIST+, SpaGE, SpaIM, stImpute and Tangram. Metrics were evaluated using per-gene SSIM, PCC and RMSE on 45 out of 50 imputed genes (the five genes are missing in the scRNA reference).

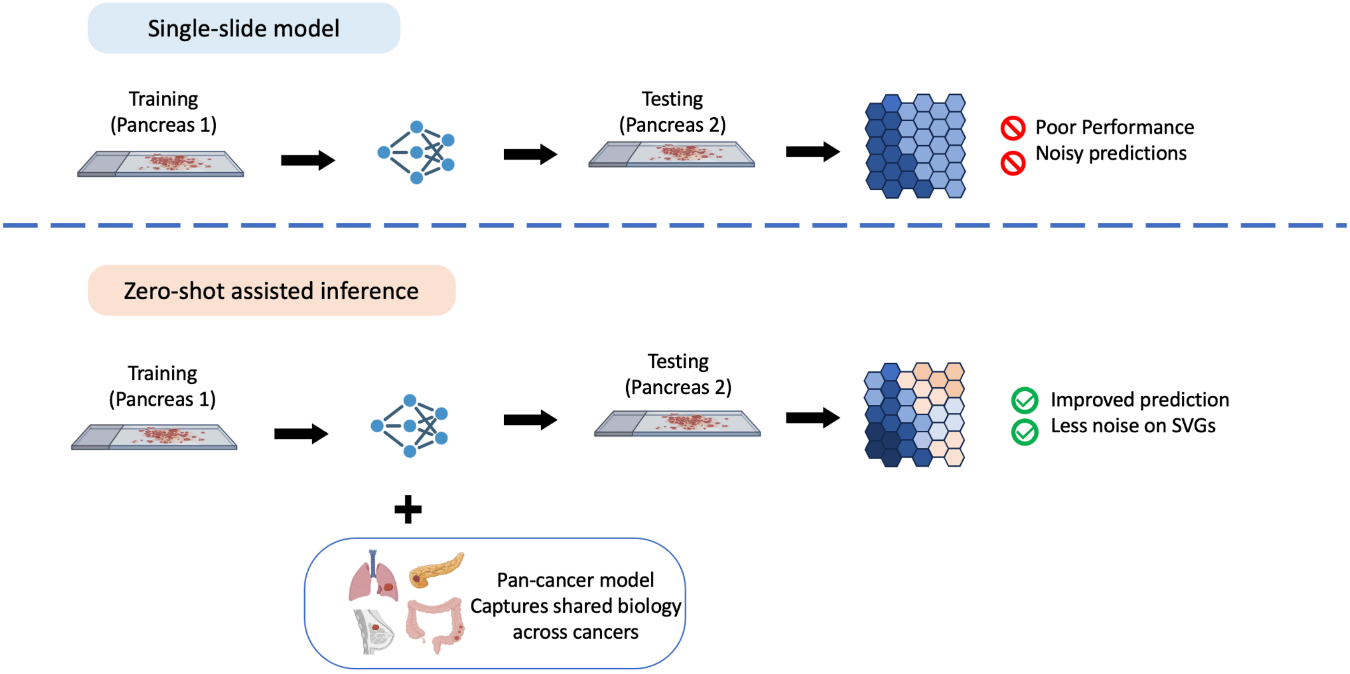


**Supplementary Figure S8:** Overview of zero-shot assisted inference. A single-slide pancreas model was augmented using a multi-cancer reference, resulting in improved spatial transcriptomic prediction and reduced noise in SVGs.

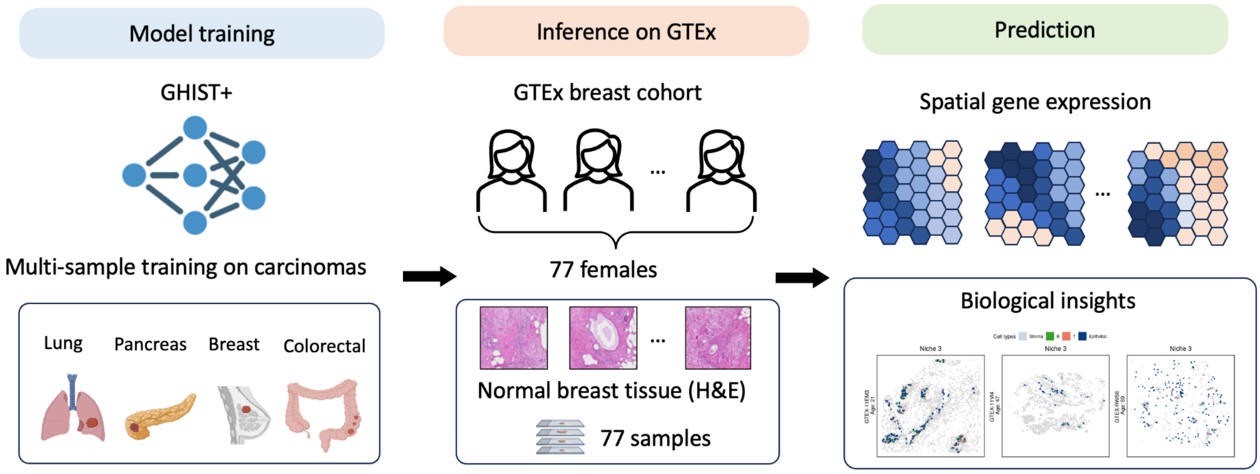


**Supplementary Figure S9:** Workflow for applying GHIST+ to normal breast tissue. A multi-sample carcinoma-trained model was used to infer spatial gene expression patterns from H&E images in the GTEx female breast cohort (77 samples), enabling downstream spatial and cellular analyses.

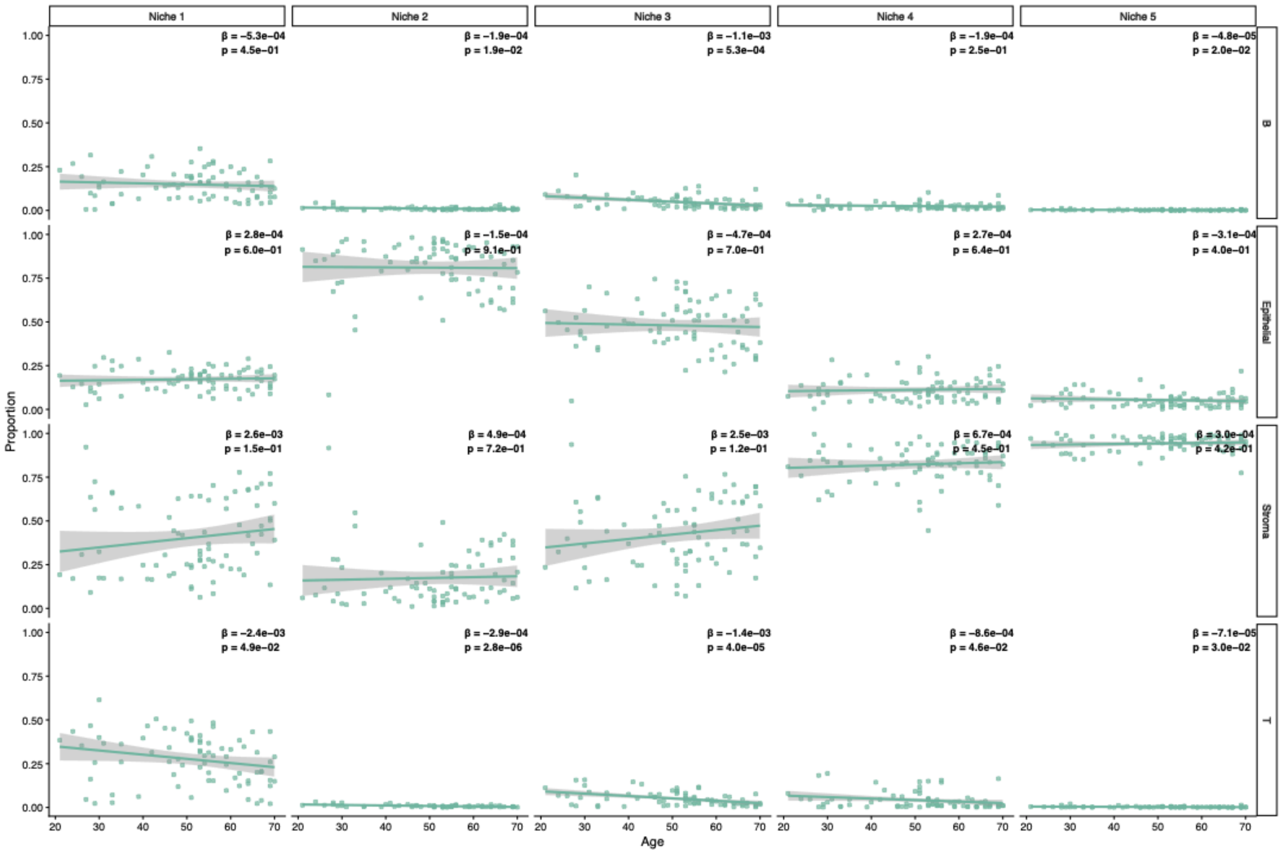


**Supplementary Figure S10: Age-associated cellular composition changes within 5 niches identified using BANKSY.** Each point represents a single sample, with B cell, endothelial cell, stromal cell and T cell proportions calculated within each niche. Regression coefficients (β) and corresponding P values are shown for each cell type within a niche.

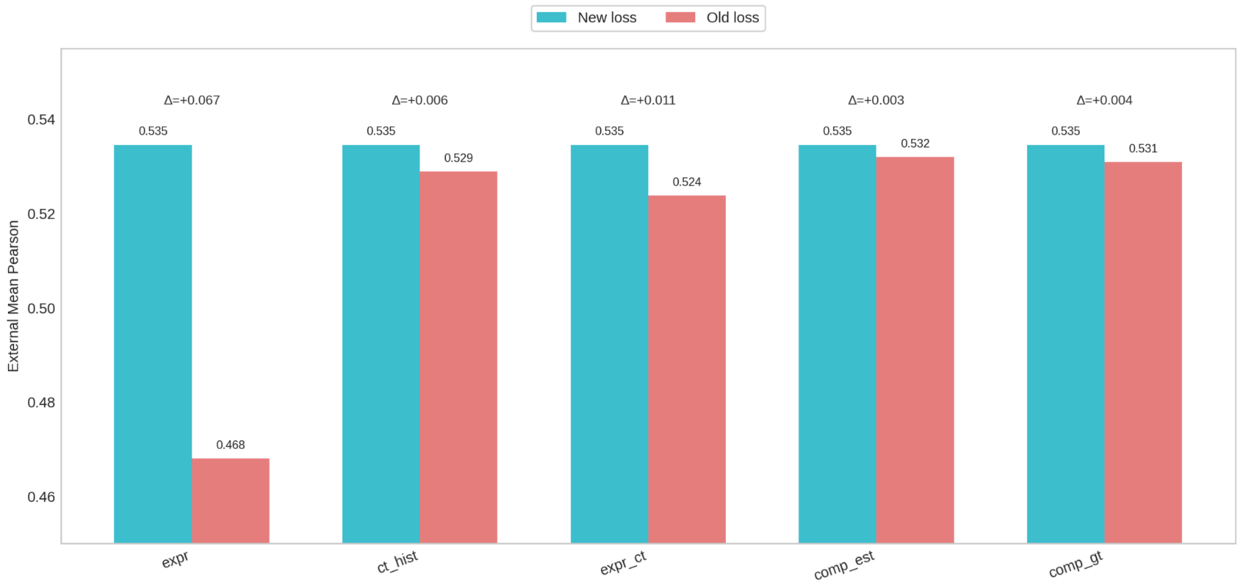


**Supplementary Figure S11: Loss ablation study**. Bar plot comparing GHIST loss function with the new GHIST+ loss functions across randomly selected patches. Losses include expression loss, cell type loss and composition loss.

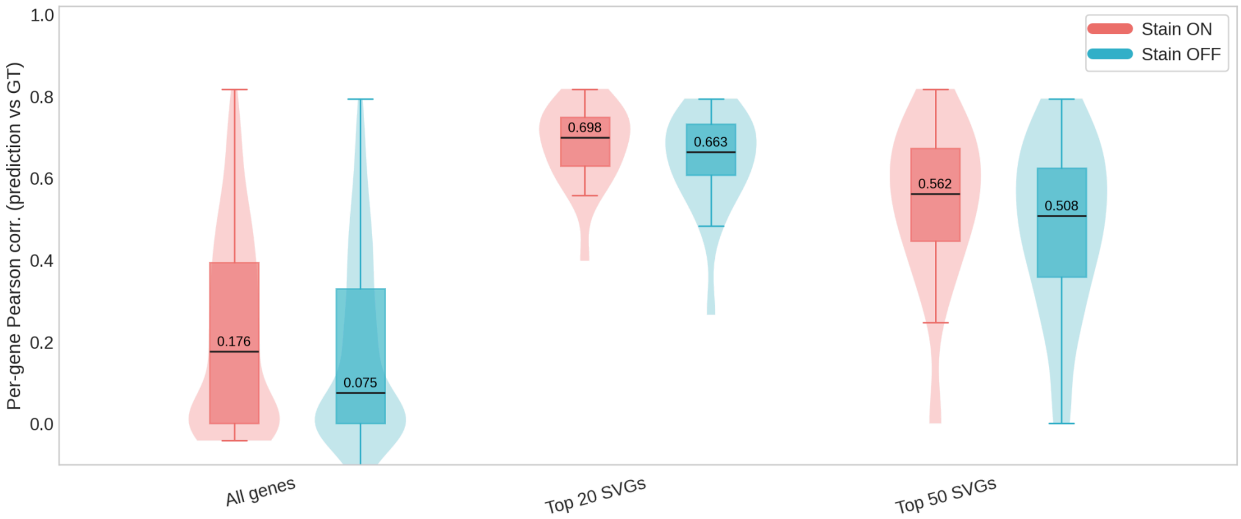


**Supplementary Figure S12: Ablation study on Macenko stain normalisation.** Stain normalisation was toggled on and off for training and testing on a randomised patch respectively. Violin boxplots show performance difference affected by stain normalisation.
